# Stability of Polygenic Scores Across Discovery Genome-Wide Association Studies

**DOI:** 10.1101/2021.06.18.449060

**Authors:** Laura M. Schultz, Alison K. Merikangas, Kosha Ruparel, Sébastien Jacquemont, David C. Glahn, Raquel E. Gur, Ran Barzilay, Laura Almasy

## Abstract

Polygenic scores (PGS) are commonly evaluated in terms of their predictive accuracy at the population level by the proportion of phenotypic variance they explain. To be useful for precision medicine applications, they also need to be evaluated at the individual patient level when phenotypes are not necessarily already known. Hence, we investigated the stability of PGS in European-American (EUR)- and African-American (AFR)-ancestry individuals from the Philadelphia Neurodevelopmental Cohort (PNC) and the Adolescent Brain Cognitive Development (ABCD) cohort using different discovery GWAS for post-traumatic stress disorder (PTSD), type-2 diabetes (T2D), and height. We found that pairs of EUR-ancestry GWAS for the same trait had genetic correlations > 0.92. However, PGS calculated from pairs of same-ancestry and different-ancestry GWAS had correlations that ranged from <0.01 to 0.74. PGS stability was higher for GWAS that explained more of the trait variance, with height PGS being more stable than PTSD or T2D PGS. Focusing on the upper end of the PGS distribution, different discovery GWAS do not consistently identify the same individuals in the upper quantiles, with the best case being 60% of individuals above the 80^th^ percentile of PGS overlapping from one height GWAS to another. The degree of overlap decreases sharply as higher quantiles, less heritable traits, and different-ancestry GWAS are considered. PGS computed from different discovery GWAS have only modest correlation at the level of the individual patient, underscoring the need to proceed cautiously with integrating PGS into precision medicine applications.

## Introduction

Polygenic scores (PGS) are increasingly being used to draw inferences regarding genetic contributions to a variety of complex anthropometric and disease-related traits. Numerous methods have been developed for computing PGS for a target population using summary statistics from a discovery genome-wide association study (GWAS) run for an independent population, with newer Bayesian-based techniques such as LDpred,^1^ SBayesR,^2^ and PRS-CS^3^ generally yielding more predictive PGS than those produced using older methodologies that rely on a combination of linkage disequilibrium (LD) clumping and *P*-value thresholding.^4^

One goal is to utilize PGS in clinical settings to facilitate the diagnosis and treatment of a wide range of heritable diseases,^5^ such as inflammatory bowel disease,^6^ diabetes,^7^ cardiovascular disease,^8;9^ cancer,^10^ Alzheimer’s disease,^11^ attention-deficit/hyperactivity disorder,^12^ major depressive disorder,^13^ bipolar disorder,^14^ and schizophrenia.^15^ While progress has been made towards reaching this goal,^16–19^ numerous challenges remain to be solved.^5;20–23^ Given that the GWAS required for computing PGS have been disproportionately run for European-ancestry populations,^24–28^ a fundamental challenge will be ensuring that diverse populations have equitable access to medically beneficial PGS,^29^ as it has been demonstrated that that PGS are less predictive when the target and discovery populations have differing genetic ancestry or varying degrees of admixture.^30–34^

Previous studies have evaluated PGS performance in terms of how well they predict phenotypes at the population level. However, if PGS are going to be adopted in the precision-medicine setting, it is also necessary to examine how well PGS perform at predicting the risk for individual patients.^35^ To this end, we examined the stability of PGS computed for individuals across discovery GWAS. Specifically, we evaluated the correlations between the PGS computed for EUR and AFR individuals from pairs of same- and different-ancestry discovery GWAS for post-traumatic stress disorder,^36;37^ type 2 diabetes,^38–40^ and height.^41;42^ These specific traits were chosen because they had sufficiently powered, publicly available AFR-ancestry GWAS. We also addressed the question of whether the same individuals were consistently identified as belonging to the top PGS quantiles. For this work, we targeted European-American (EUR) and African-American (AFR) youth from the Philadelphia Neurodevelopmental Cohort (PNC) and the Adolescent Brain Cognitive Development Study (ABCD).

## Subjects and Methods

### Philadelphia Neurodevelopmental Cohort (PNC)

Genotype data for the PNC, a population-based sample of youth who were ages 8-21 at the time of study enrollment,^43^ were obtained from dbGaP (phs000607.v2.p2). Biological samples from PNC subjects were genotyped in fifteen batches (Table S1) using ten different types of Affymetrix and Illumina arrays by the Center for Applied Genomics at the Children’s Hospital of Philadelphia.^44^ Analysis was limited to the 5,239 EUR and 3,260 AFR ancestry individuals for whom genotype data were available after the quality-control (QC) process described below.

### Adolescent Brain Cognitive Development Study (ABCD)

Results were replicated using post-QC genotype data for 5,815 EUR and 1,741 AFR individuals in the independent ABCD cohort (NDA #2573, fix release 2.0.1). This cohort is comprised of adolescents who were ages 9-10 at the time that their saliva samples were collected for genotyping.^45^ The Rutgers University Cell and DNA Repository stored and genotyped all samples using the Affymetrix NIDA SmokeScreen array.

### Quality Control and Imputation

The PNC dataset was processed by array batch and merged after imputation, whereas the ABCD dataset was processed as a single batch. For each batch, PLINK 1.9^46^ was used to remove single nucleotide polymorphisms (SNPs) with > 5% missingness, samples with more than 10% missingness, and samples with a genotyped sex that did not match the reported sex phenotype. As a final step, each batch was checked with a pre-imputation perl script that compared SNP frequencies against the 1000 Genomes ALL reference panel.^47^ This script fixed strand reversals and improper Ref/Alt assignments and also removed palindromic A/T and C/G SNPs with minor allele frequency (MAF) > 0.4, SNPs with alleles that did not match the reference panel, SNPs with allele frequencies differing by more than 0.2 from the reference, and SNPs not present in the reference panel.

Genotypes were phased (Eagle v.2.4) and imputed by chromosome to the 1000 Genomes Other/Mixed GRCh37/hg19 reference panel (Phase 3 v.5) using Minimac 4 via the Michigan Imputation Server.^48^ The fifteen imputation batches for the PNC dataset were merged by chromosome, and then post-imputation QC was run on the merged chromosome files using bcftools.^49^ Only polymorphic sites with imputation quality *R*^2^ ≥ 0.7 and MAF ≥ 0.01 were included in the final PLINK 1.9 hard-call PNC and ABCD post-imputation datasets.

### Ancestry and Kinship Analysis

Multi-dimensional scaling (MDS) was conducted using KING (v.2.2.4)^50^ to identify the top ten ancestry principal components (PCs) for each sample. These PCs were projected onto the 1000 Genomes PC space, and genetic ancestry was inferred using the e1071^51^ support vector machines package in R^52^ (Figure 1). Based on these inferences, AFR- and EUR-ancestry cohorts were created for the PNC and ABCD datasets; all other ancestry groups were excluded from further analysis. A second round of unprojected MDS was then performed within the EUR- and AFR-ancestry groups to produce ten PCs that were regressed out of the standardized PGS to adjust for array batch effects and genetic ancestry (Figures S1-S5).

**Figure 1.**
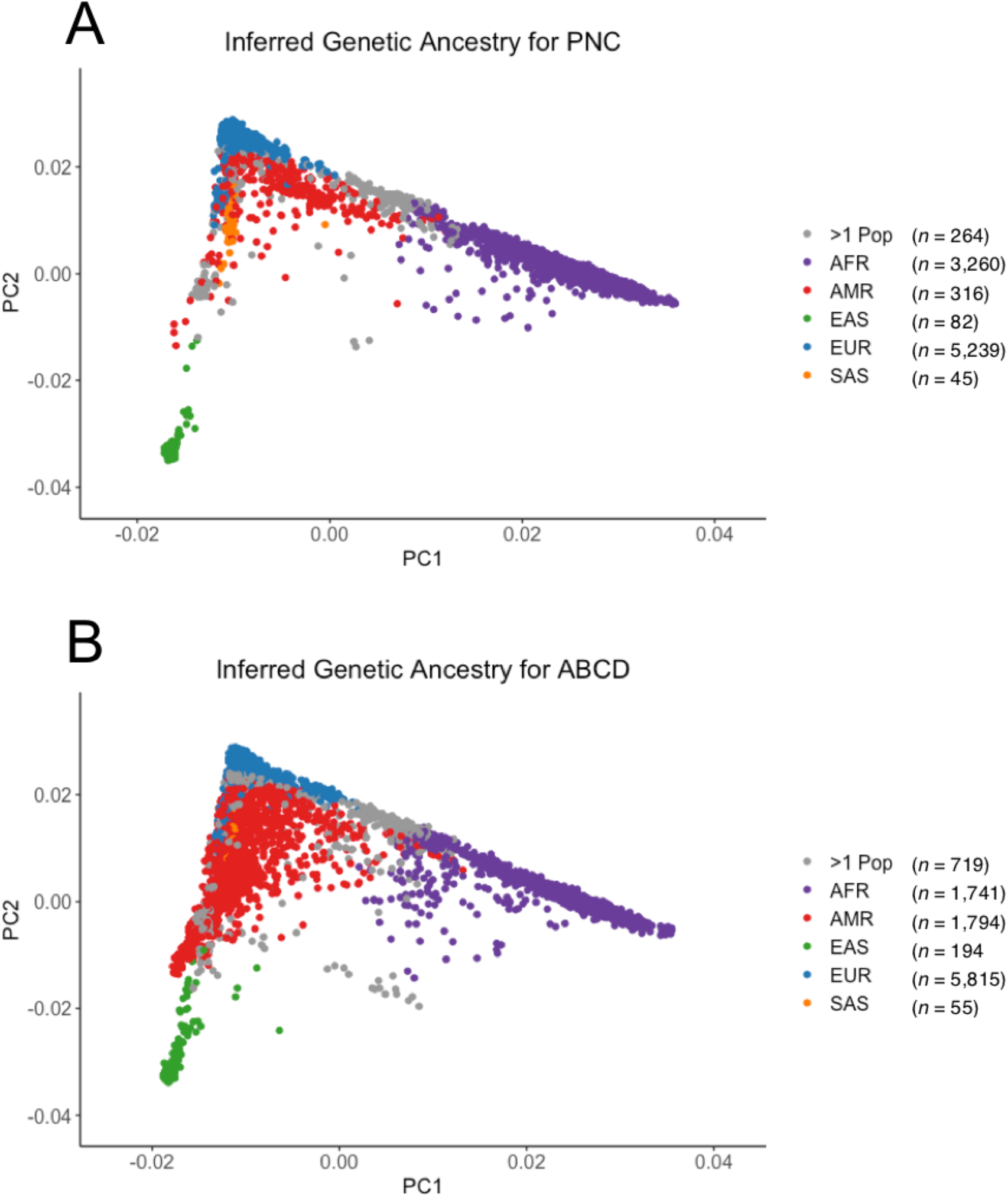
First and second principal components (PCs) of cohort genotypes. PCs were computed and projected to a 1000 Genomes reference using KING.^50^ Colors indicate inferred genetic ancestry for the (A) 9,206 Philadelphia Neurodevelopmental Cohort (PNC) and (B) 10,318 Adolescent Brain Cognitive Development (ABCD) genotyped samples.

KING was also used to identify all pairwise relationships out to third degree relatives based on estimated kinship coefficients and inferred IBD segments. Although the PNC was not recruited as a family study, it does include some related individuals (i.e., siblings and cousins). We ran a sensitivity analysis using a reduced PNC dataset that included only one individual from each family (chosen as the lowest individual ID number for a given family ID number), which reduced the size of the PNC EUR cohort from 5,239 to 4,928 and the AFR cohort from 3,260 to 2,954. After establishing that the PNC PTSD PGS correlation results obtained using only unrelated individuals did not differ meaningfully from those obtained using the full dataset (Tables S4 and S5), we performed all subsequent analyses using the complete EUR and AFR cohorts.

### Polygenic Score Computation with PRS-CS

PRS-CS^3^ was used to infer posterior effects by chromosome for the SNPs in a given dataset that overlapped with both the discovery GWAS summary statistics and an external 1000 Genomes LD panel that was matched to the ancestry group used for the discovery GWAS. Posterior effects were only inferred for SNPs located on the 22 autosomal chromosomes. PGS for the EUR and AFR subsets of PNC and ABCD were computed using both EUR and AFR discovery GWAS for post-traumatic stress disorder (PTSD),^36;37^ Type-2 diabetes (T2D),^38–40^ and height^41;42^ (Table 1). To ensure convergence of the underlying Gibbs sampler algorithm, we ran 25,000 Markov chain Monte Carlo (MCMC) iterations and designated the first 10,000 MCMC iterations as burn-in. The PRS-CS global shrinkage parameter was set to 0.01 when the discovery GWAS had a SNP sample size that was less than 200,000; otherwise, it was learned from the data using a fully Bayesian approach. Default settings were used for all other PRS-CS parameters. Given the stochastic nature of the Bayesian algorithm used by PRS-CS, PGS replicability was confirmed by completing multiple PRS-CS runs using the same discovery GWAS. Raw PGS were produced from the posterior effects using PLINK 1.9. R^52^ was used to standardize the PGS for a given cohort to mean = 0 and SD = 1. Standardized PGS were then adjusted by regressing out the first ten within-ancestry PCs.

**Table 1.** Discovery GWAS used to compute polygenic scores with PRS-CS.

| Trait | Discovery GWAS | GWAS Ancestry | SNP Sample Size<sup>a</sup> (for PRS-CS) | SNP Count<sup>b</sup> for PNC PGS Calculations | SNP Count for ABCD PGS Calculations |
| --- | --- | --- | --- | --- | --- |
| PTSD | Nievergelt et al. (2019)<sup>37</sup><br>(Freeze 2 PGC) | AFR | 11,321 | 1,162,502 | 1,064,574 |
|  |  | EUR | 70,237 | 1,087,435 | 1,016,161 |
|  | Duncan et al. (2018)<sup>36</sup><br>(Freeze 1 PGC) | AFR | 9,691 | 1,157,302 | 1,059,197 |
|  |  | EUR | 9,954 | 1,086,644 | 1,015,369 |
| T2D | Chen et al. (2019)<sup>40</sup> | AFR | 4,146 | 1,114,936 | 1,020,579 |
|  | Scott et al. (2017)<sup>38</sup><br>(DIAGRAM) | EUR | 152,599 | 1,087,724 | 1,016,440 |
|  | Mahajan et al. (2018)<sup>39</sup><br>(DIAGRAM) | EUR | 231,420 | 1,089,613 | 1,018,372 |
| Height | Marouli et al. (2017)<sup>42</sup><br>(GIANT) | AFR | 27,494 | 18,580 | 15,720 |
|  |  | EUR | 381,625 | 18,035 | 15,767 |
|  | Wood et al. (2014)<sup>41</sup><br>(GIANT) | EUR | 252,048 | 987,760 | 920,889 |

PTSD, post-traumatic stress disorder; T2D, type 2 diabetes; PNC, Philadelphia Neurodevelopmental Cohort; PGS, polygenic score; ABCD, Adolescent Brain Cognitive Development Study; PGC, Psychiatric Genomics Consortium; DIAGRAM, Diabetes Genetics Replication and Meta-Analysis Consortium; GIANT, Genetic Investigation of Anthropometric Traits Consortium.
<sup>a</sup>PRS-CS requires a single SNP sample size; see Supplemental Methods for details.
<sup>b</sup>The "SNP count" is the number of SNPs in common between the discovery GWAS, the PRS-CS LD panel, and the genomic dataset.

### SNP Heritability Estimation with LDSC

LD score regression (LDSC)^53; 54^ was used to calculate SNP-based heritability 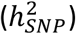 on the observed scale for each EUR-ancestry discovery GWAS that we used to generate PGS. We also calculated 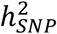 on the liability scale for PTSD and T2D by incorporating the sample prevalence and estimated population prevalence into the calculation (Table S14). The standard error associated with a given 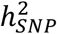 calculation was estimated using a block jackknife over blocks of adjacent SNPs. Given that LDSC may yield biased estimates for admixed populations^55^ and that we only had access to GWAS summary statistics, we did not calculate SNP heritability for the AFR-ancestry discovery GWAS.

### Statistical Analysis

All statistics and graphical displays were generated using R.^52^ Pearson correlation coefficients were calculated to assess the strength of correlations between PC-adjusted standardized PGS that were calculated for a given trait using different discovery GWAS. Statistical significance was determined with two-tailed *t*-tests for linear association. For each comparison, counts were also made of the number of samples in common at or above the 80^th^ percentile, the 90^th^ percentile, and the 95^th^ percentile of the adjusted standardized PGS distributions.

## Results

### Reproducibility of Bayesian Posterior Effects

Given that PRS-CS relies on Bayesian methodology to infer posterior effects for the SNPs on each chromosome,^3^ it was necessary to confirm that we had used enough Markov chain Monte Carlo (MCMC) iterations and burn-in trials to ensure convergence of the underlying Gibbs sampler algorithm. We checked for convergence indirectly by assessing the correlation between the posterior effects calculated across multiple runs for a given chromosome (Figure 2). The PRS-CS default setting of 1000 MCMC iterations with the first 500 iterations serving as burn-in produced relatively inconsistent posterior effects (*r* ≈ 0.8), suggesting incomplete convergence. The correlation between the posterior effects computed during multiple runs of PRS-CS improved to *r* ≈ 0.98 when we increased the number of MCMC iterations to 10,000 (5,000 burn-in) and further improved to *r* > 0.99 for both large and small chromosomes when we used 25,000 MCMC iterations (10,000 burn-in). Given that the computational time increases substantially as more MCMC iterations are run, we opted to use 25,000 MCMC iterations with the first 10,000 as burn-in rather than pursuing even stronger correlations.

**Figure 2.**
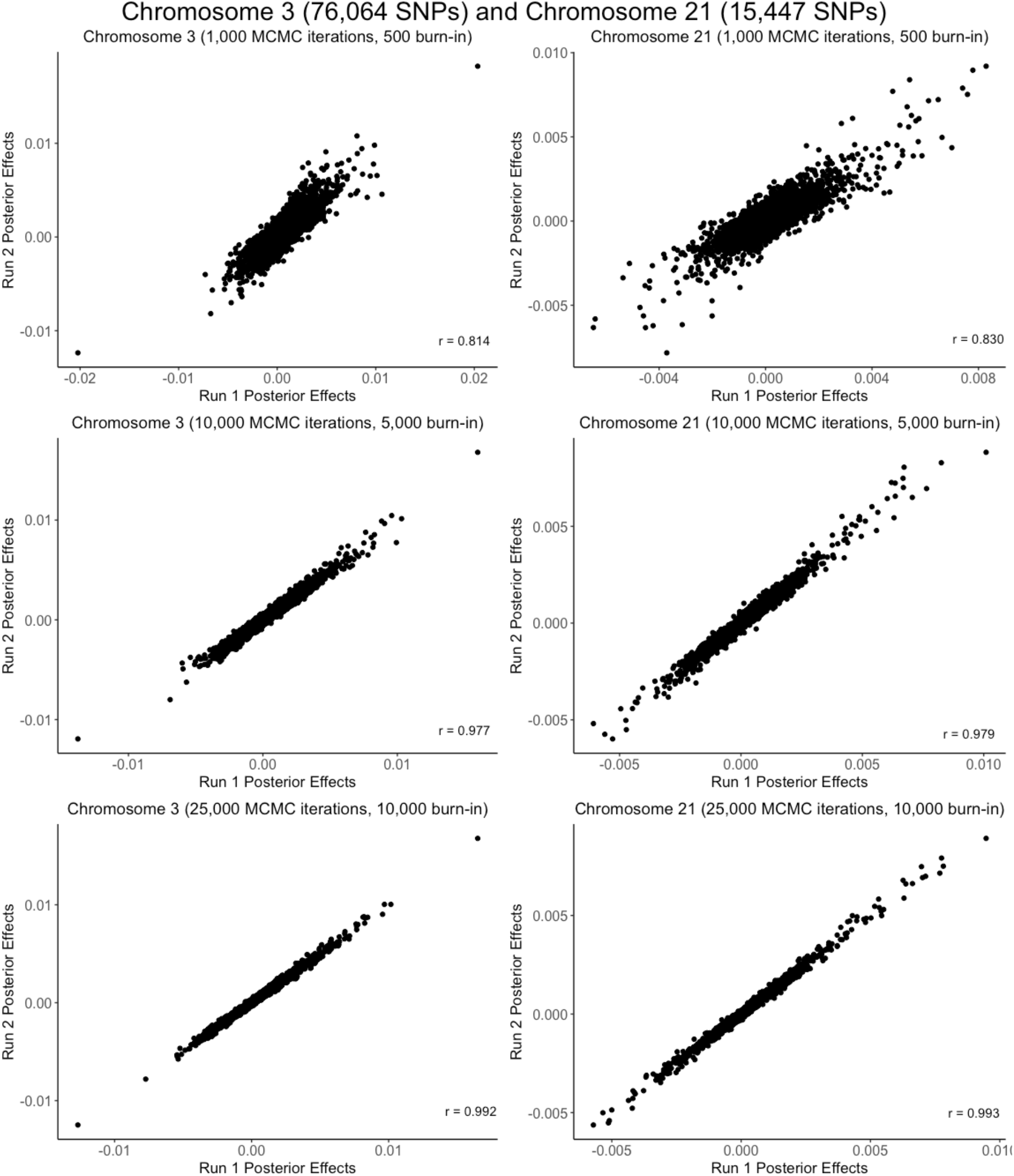
Reproducibility of Bayesian posterior effects computed by PRS-CS. As illustrated for chromosome 3 (76,064 SNPs) and chromosome 21 (15,447 SNPs) using the Nievergelt et al. (2019)^37^ EUR PTSD discovery GWAS with the PNC EUR dataset, posterior effects were more strongly correlated between PRS-CS runs as the number of MCMC iterations (and burn-in iterations) increased.

### Reproducibility of PGS Computed from the Same Discovery GWAS

The next concern was whether the PGS calculated by PLINK 1.9 from the Bayesian posterior effects would also be reproducible across PRS-CS runs. To address this question, we ran PRS-CS twice using the Psychiatric Genomics Consortium (PGC) Freeze 2 PTSD discovery GWAS^37^, and calculated PGS from both sets of posterior effects. For both the EUR and AFR PNC cohorts, the correlation between the adjusted PGS was greater than 0.999 (Figure 3). Hence, we are confident that PRS-CS yields reproducible PGS for a given discovery GWAS provided that enough MCMC iterations are used.

**Figure 3.**
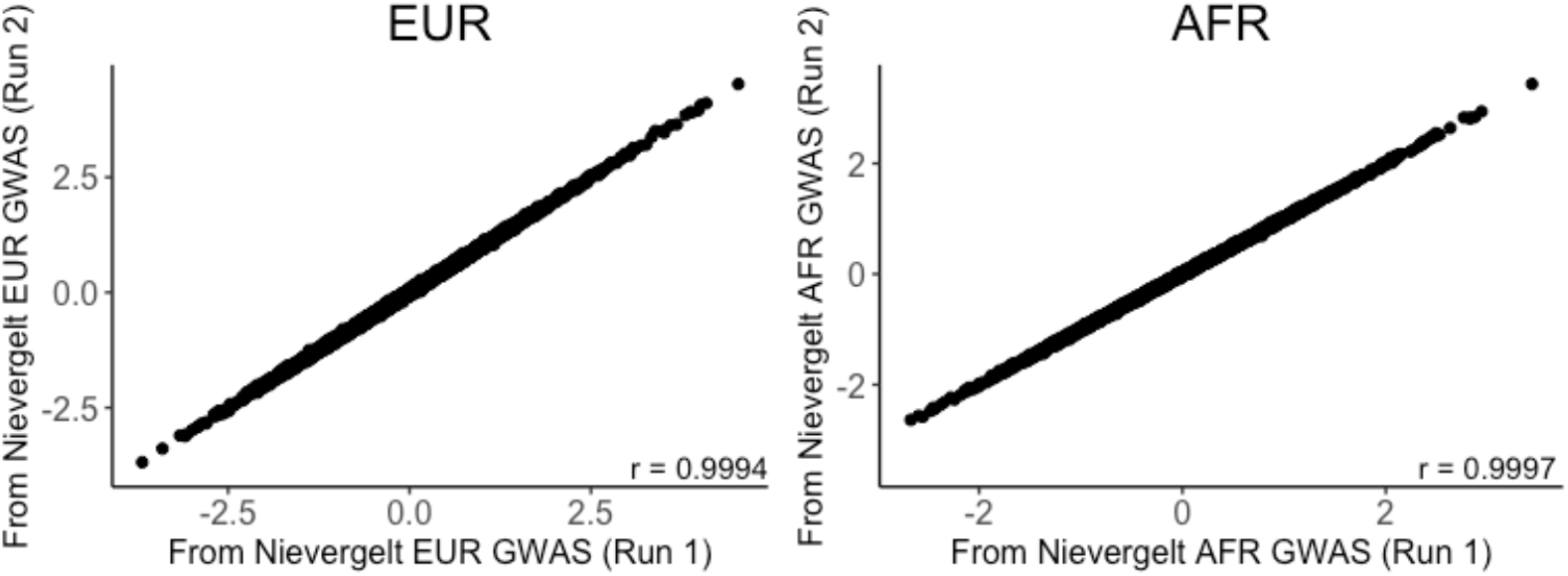
Reproducibility of PGS across multiple runs of PRS-CS. PC-adjusted standardized PGS computed from posterior effects generated by two runs of PRS-CS using the same PTSD discovery GWAS from Nievergelt et al. (2019)^37^ had correlations greater than *r* = 0.999 for both the EUR (*n* = 5239) and AFR (*n* = 3260) cohorts of PNC.

### Stability of PGS Computed from Different Same-Ancestry Discovery GWAS

Of the three traits that we analyzed, only PTSD had two publicly available AFR-ancestry GWAS.^36;37^ We computed PGS using both GWAS for each AFR-ancestry individual and then assessed the correlation between the two sets of PGS (Figure 4). We found a moderately strong positive correlation between the PGS computed from the PGC Freeze 1^36^ and Freeze 2^37^ AFR-ancestry PTSD GWAS for the AFR-ancestry cohorts of both PNC (*r* = 0.696, *t*(3258) = 55.26, *P* < 2 × 10^−16^) and ABCD (*r* = 0.657, *t*(1739) = 36.34, *P* < 2 × 10^−16^).

**Figure 4.**
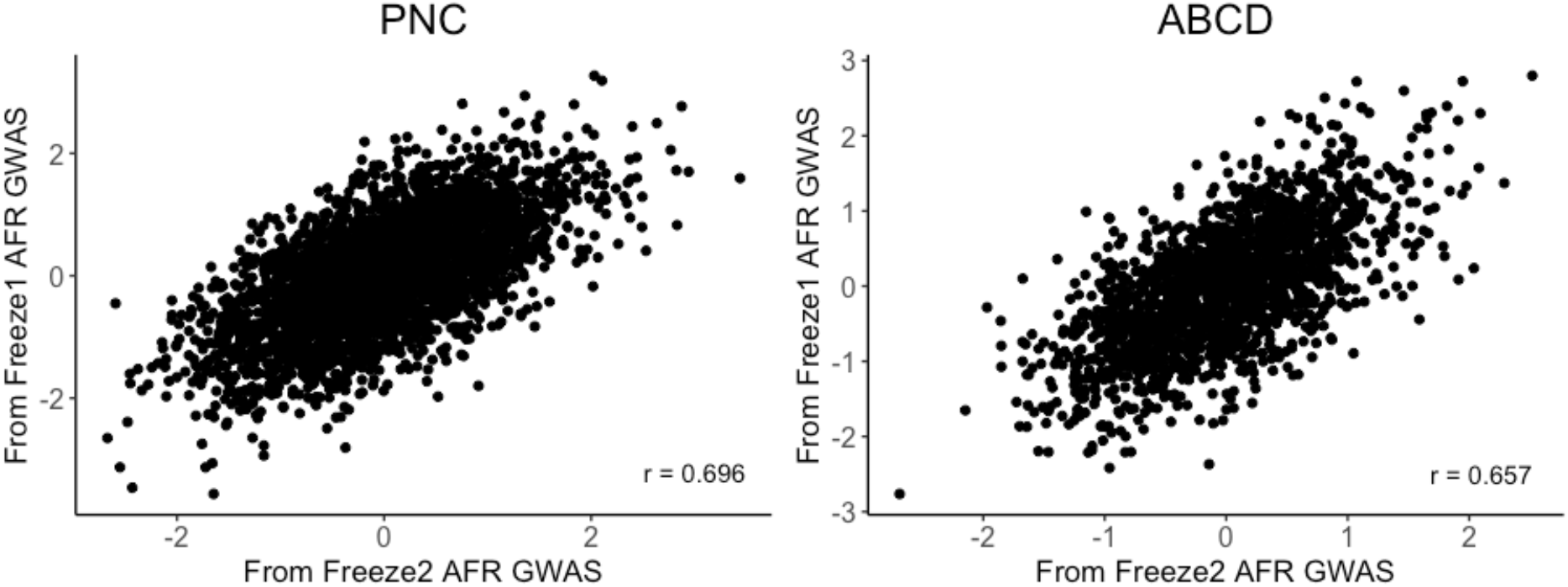
Correlation between PGS computed from two different AFR-ancestry PTSD discovery GWAS for AFR-ancestry individuals. Significant positive correlations were observed between the AFR PGS computed from the PGC Freeze1^36^ and Freeze2^37^ AFR PTSD GWAS for both the PNC (*r* = 0.696, *t*(3258) = 55.26, *P* < 2 × 10^−16^) and ABCD (*r* = 0.657, *t*(1739) = 36.34, *P* < 2 × 10^−16^) AFR cohorts.

The wider availability of EUR-ancestry GWAS allowed us to compute PGS for EUR-ancestry individuals using pairs of EUR-ancestry discovery GWAS for PTSD,^36;37^ T2D,^38;39^ and height^41;42^ (Figure 5). Statistically significant positive correlations between the pairs of PGS were observed for all three traits for both the PNC (Table S8) and ABCD (Table S9) EUR-ancestry cohorts, with the strongest association observed between the height PGS (PNC: *r* = 0.736; ABCD: *r* = 0.734) and the weakest observed for the PTSD PGS (PNC: *r* = 0.392; ABCD: *r* = 0.378).

**Figure 5.**
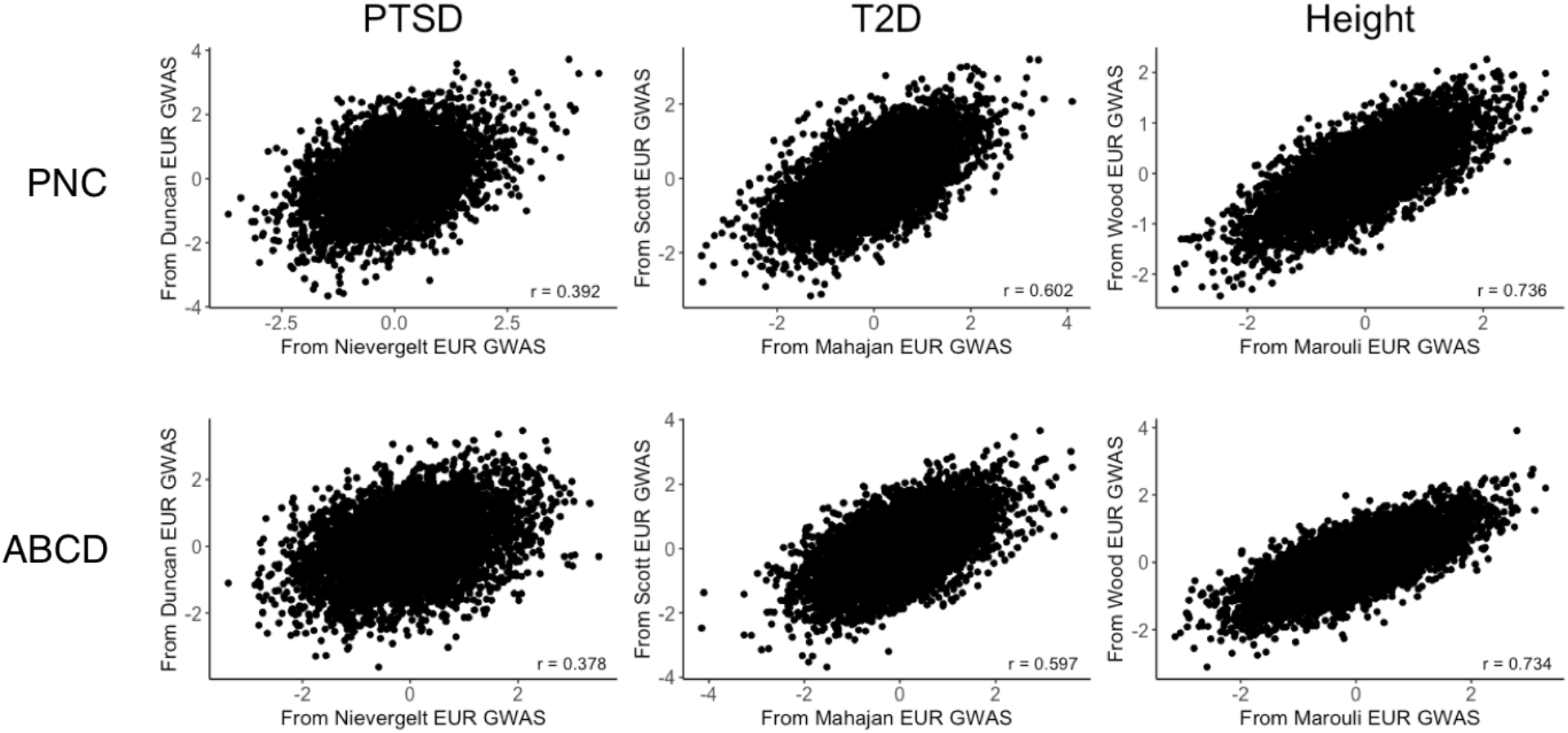
Correlation between PGS computed from two different EUR-ancestry discovery GWAS for EUR-ancestry individuals. Pairs of PGS computed for the EUR samples of PNC (*n* = 5239) and ABCD (*n* = 5815) using two different EUR discovery GWAS for PTSD,^36; 37^ T2D,^38; 39^ and height^41;42^ all showed significant positive correlations.

### Stability of PGS Computed from Different-Ancestry Discovery GWAS

Given the scarcity of AFR-ancestry GWAS, it is often tempting to compute PGS for AFR-ancestry individuals using EUR-ancestry discovery GWAS. To assess the feasibility of this approach, we computed PGS for AFR-ancestry individuals in PNC and ABCD using both AFR-ancestry discovery GWAS and EUR-ancestry GWAS and then assessed the correlation between the two sets of PGS (Figure 6).

**Figure 6.**
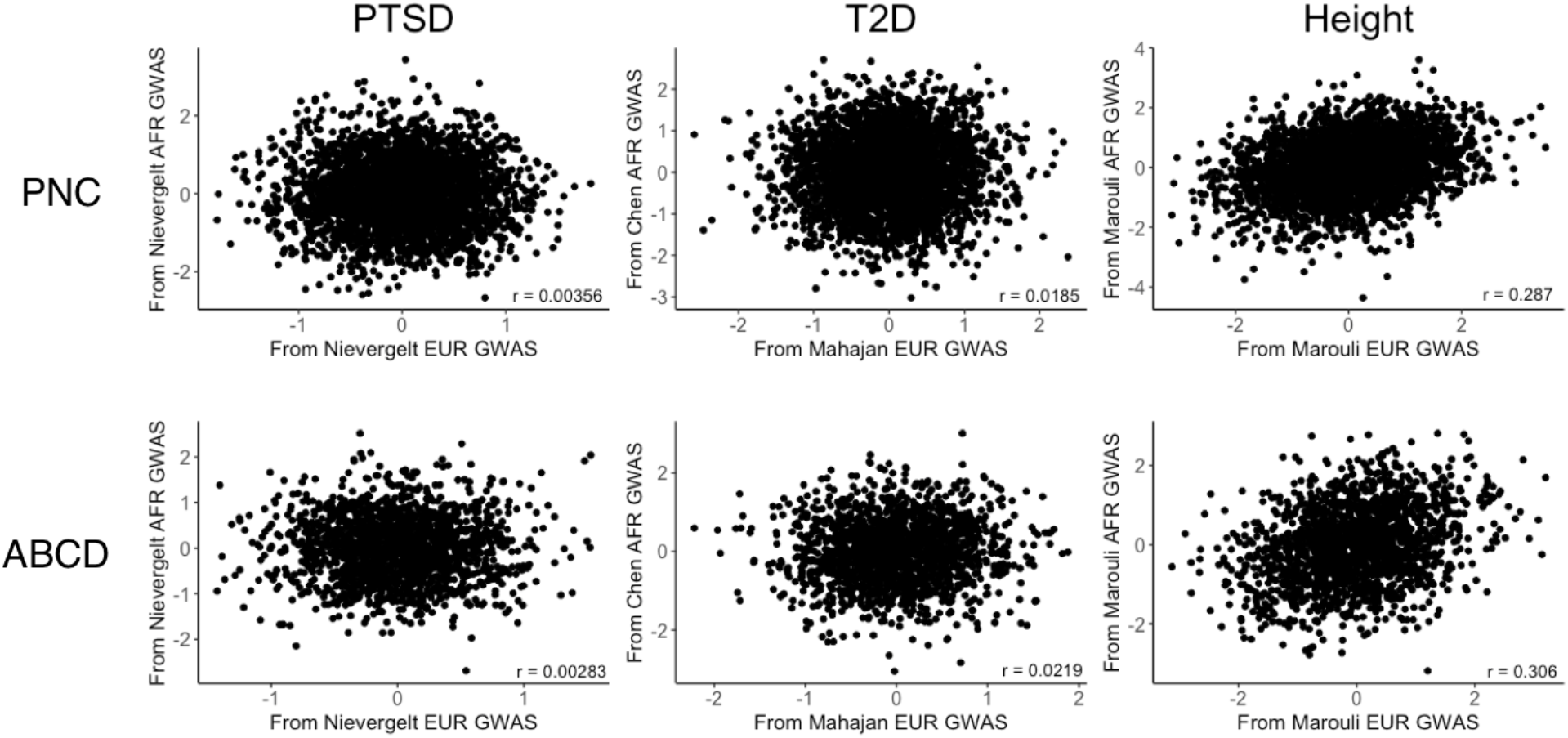
Correlation between PGS computed from AFR-ancestry and EUR-ancestry discovery GWAS for AFR-ancestry individuals. Pairs of PGS computed for the AFR samples of PNC and ABCD from the newer EUR and AFR discovery GWAS were not significantly correlated for either PTSD^36;37^ or T2D,^39; 40^ but there was a significant positive correlation for height.^42^

For PTSD, there was no significant correlation between the PGS computed from the newer Freeze 2 PGC AFR and EUR discovery GWAS^37^ for AFR-ancestry individuals in either PNC (*r* = 0.00356, *t*(3258) = 0.203, *P* = 0.839) or ABCD (*r* = 0.00283, *t*(1739) = 0.118, *P* = 0.906). The AFR PGS computed using the Freeze 1 PGC PTSD AFR and EUR discovery GWAS^36^ were uncorrelated for ABCD (*r* = −0.00320, *t*(1739) = −0.133, *P* = 0.894), but we observed a weak positive correlation for PNC (*r* = 0.0417, *t*(3258) = 2.379, *P* = 0.0174).

We made the same different-ancestry GWAS comparisons for the EUR-ancestry individuals in the PNC and ABCD study populations (Figure 7). As was the case for AFR-ancestry individuals, we found no significant correlation between PGS computed from the PGC Freeze 2 EUR- and AFR-ancestry PTSD discovery GWAS.^37^ While we observed no significant correlation between the PGS computed using the PGC Freeze 1 EUR- and AFR-ancestry PTSD discovery GWAS for EUR-ancestry individuals in ABCD (*r* = −0.00109, *t*(5813), *P* = 0.934), we did observe a weak positive correlation for the EUR cohort of PNC (*r* = 0.0379, *t*(5237) = 2.746, *P* = 0.0065).

**Figure 7.**
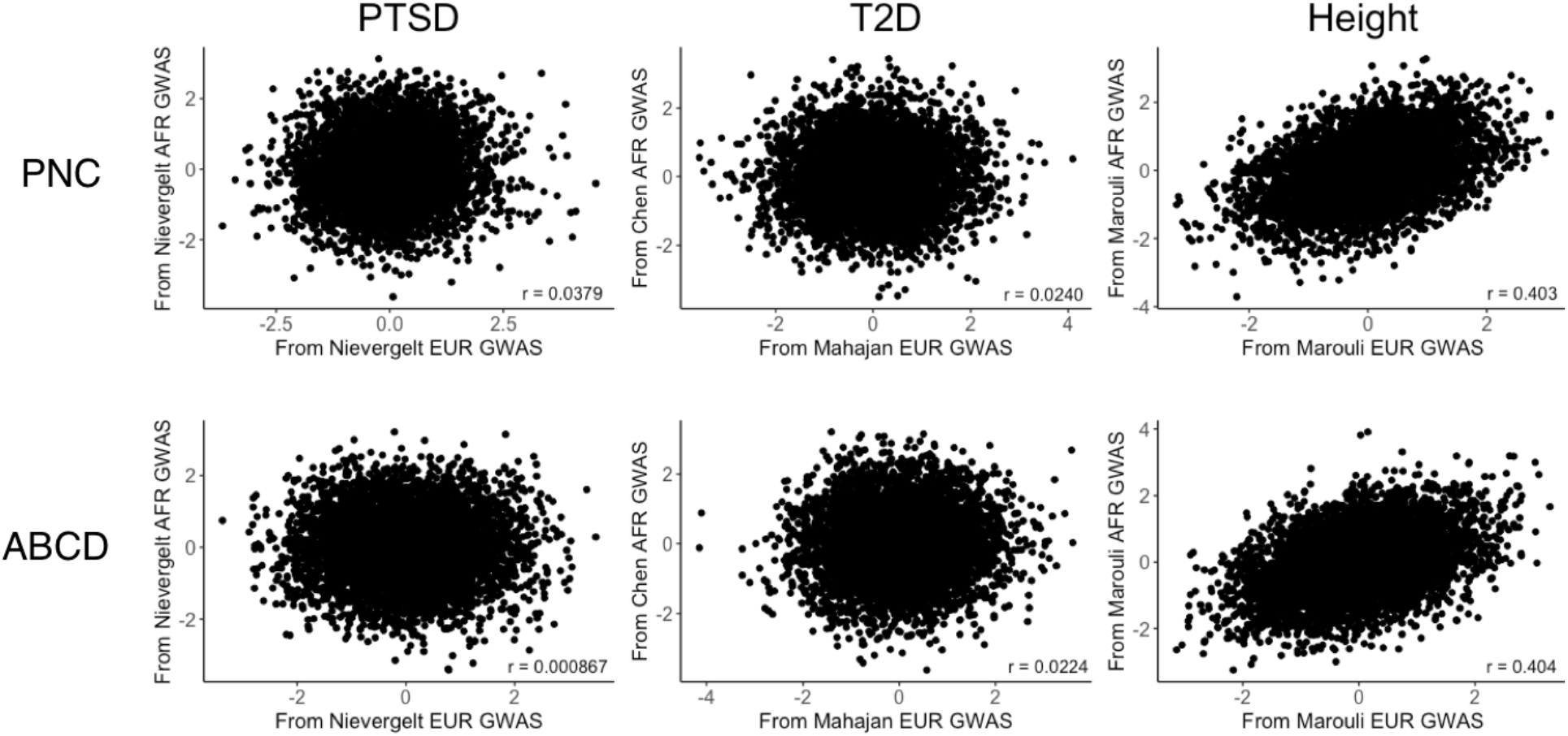
Correlation between PGS computed from EUR-ancestry and AFR-ancestry discovery GWAS for EUR-ancestry individuals. Pairs of PGS computed for the EUR samples of PNC and ABCD from the newer EUR and AFR discovery GWAS were not significantly correlated for either PTSD^36; 37^ or T2D,^39; 40^ but there was a significant positive correlation for height.^42^

We compared T2D PGS computed from an AFR-ancestry discovery GWAS^40^ to those computed using two EUR discovery GWAS^38;39^ published by the Diabetes Genetics Replication and Meta-Analysis (DIAGRAM) consortium. The newer EUR-ancestry T2D discovery GWAS^39^ yielded PGS that were uncorrelated with those computed from the AFR-ancestry discovery GWAS^40^ for the AFR-ancestry individuals in both PNC (*r* = 0.0185, *t*(3258) = 1.055, *P* = 0.292) and ABCD (*r* = 0.0219, *t*(1739) = 0.912, *P* = 0.362). Similarly, there was no significant correlation between the different-ancestry T2D PGS that we computed for the EUR-ancestry individuals in PNC (*r* = 0.0240, *t*(5237) = 1.739, *P* = 0.082) and ABCD (*r* = 0.0224, *t*(5813) = 1.71, *P* = 0.0872). We observed a weak positive correlation between the PGS computed from the older EUR-ancestry T2D discovery GWAS^38^ and the PGS computed from the AFR-ancestry T2D discovery GWAS^40^ for the PNC AFR cohort (*r* = 0.0432, *t*(3258) = 2.469, *P* = 0.0136), but there were no significant correlations between the two sets of PGS computed for the ABCD AFR cohort (*r* = − 0.0458, *t*(1739) = −1.911, *P* = 0.0562), the PNC EUR cohort (*r* = 0.00528, *t*(5237) = 0.382, *P* = 0.703), or the ABCD EUR cohort (*r*= 0.0188, *t*(5813) = 1.431, *P* = 0.152).

We also computed different-ancestry PGS using EUR- and AFR-ancestry height discovery GWAS that we obtained from the Genetic Investigation of Anthropometric Traits (GIANT) consortium.^41;42^ We observed significant positive correlations between the PGS computed from the newer EUR- and AFR-ancestry height discovery GWAS^42^ for the PNC AFR (*r* = 0.287, *t*(3258) = 17.09, *P* < 2 × 10^−16^), ABCD AFR (*r* = 0.306, *t*(1739) = 13.42, *P* < 2 × 10^−16^), PNC EUR (*r* = 0.403, *t*(5237) = 31.82, *P* < 2 × 10^−16^), and ABCD EUR (*r* = 0.404, *t*(5813) = 33.69, *P* < 2 × 10^−16^) cohorts. Likewise, we found significant positive correlations between the PGS computed from the older EUR-ancestry height discovery GWAS^41^ and the AFR-ancestry height discovery GWAS^42^ for the PNC AFR (*r* = 0.258, *t*(3258) = 15.22, *P* < 2 × 10^−16^), ABCD AFR (*r* = 0.312, *t*(1739) = 13.68, *P* < 2 < 10^−16^), PNC EUR(*r* = 0.335, *t*(5239) = 25.25, *P* < 2 × 10^−16^), and ABCD EUR (*r* = 0.327, *t*(5813)= 26.39, *P* < 2 × 10^−16^) cohorts. As was the case for T2D, there was only one AFR-ancestry height discovery GWAS^42^ available to use for computing PGS.

The supplement includes complete statistical results for the comparisons between PGS computed from different discovery GWAS for the PNC AFR (Table S6), ABCD AFR (Table S7), PNC EUR (Table S8), and ABCD EUR (Table S9) cohorts.

### Quantile-Based Comparisons

Given that there was little to no correlation between PGS computed from different discovery GWAS, we considered the possibility that there would be more stability if we focused on the individuals who had PGS located in the upper tail of the distribution, as those are the individuals who would presumably be most at risk for a disease trait. Considering the top 20%, 10%, and 5% of PC-adjusted standardized PGS, we counted how many individuals were jointly identified as being at or above a given percentile of the PGS computed from two different discovery GWAS. See the supplement for complete results of these analyses for the PNC AFR (Table S10), ABCD AFR (Table S11), PNC EUR (Table S12), and ABCD EUR (Table S13) cohorts.

As a baseline comparison, we determined the degree of overlap between the individuals in the top quantiles of PGS computed from two PRS-CS runs using the Freeze 2 AFR- and EUR-ancestry PTSD discovery GWAS^37^ for the AFR (Figure 8A) and EUR (Figure 9A) PNC cohorts, respectively. Of the *n* = 3260 individuals in the PNC AFR cohort, there are *n* = 652 individuals with PGS at or above the 80^th^ percentile, *n* = 326 with PGS at or above the 90^th^ percentile, and *n* = 163 with PGS at or above the 95^th^ percentile of PGS. We found an overlap of 644 of the 652 AFR-ancestry individuals who had PGS at or above the 80^th^ percentile from the two runs using the same AFR-ancestry PTSD discovery GWAS, which is a 98.7% overlap. Comparable degrees of overlap were observed between the PNC AFR-ancestry individuals with PTSD PGS at or above the 90^th^ (318/326 = 0.975) and 95^th^ (161/163 = 0.988) percentiles. Similarly, the proportional overlap between the PTSD PGS computed from two PRS-CS runs using the Freeze 2 EUR-ancestry PTSD discovery GWAS^37^ for the EUR-ancestry cohort (*n* = 5239) was 1026/1048 = 0.979 at or above the 80^th^ percentile, 513/524 = 0.979 at or above the 90^th^ percentile, and 255/262 = 0.973 at or above the 95^th^ percentile.

**Figure 8.**
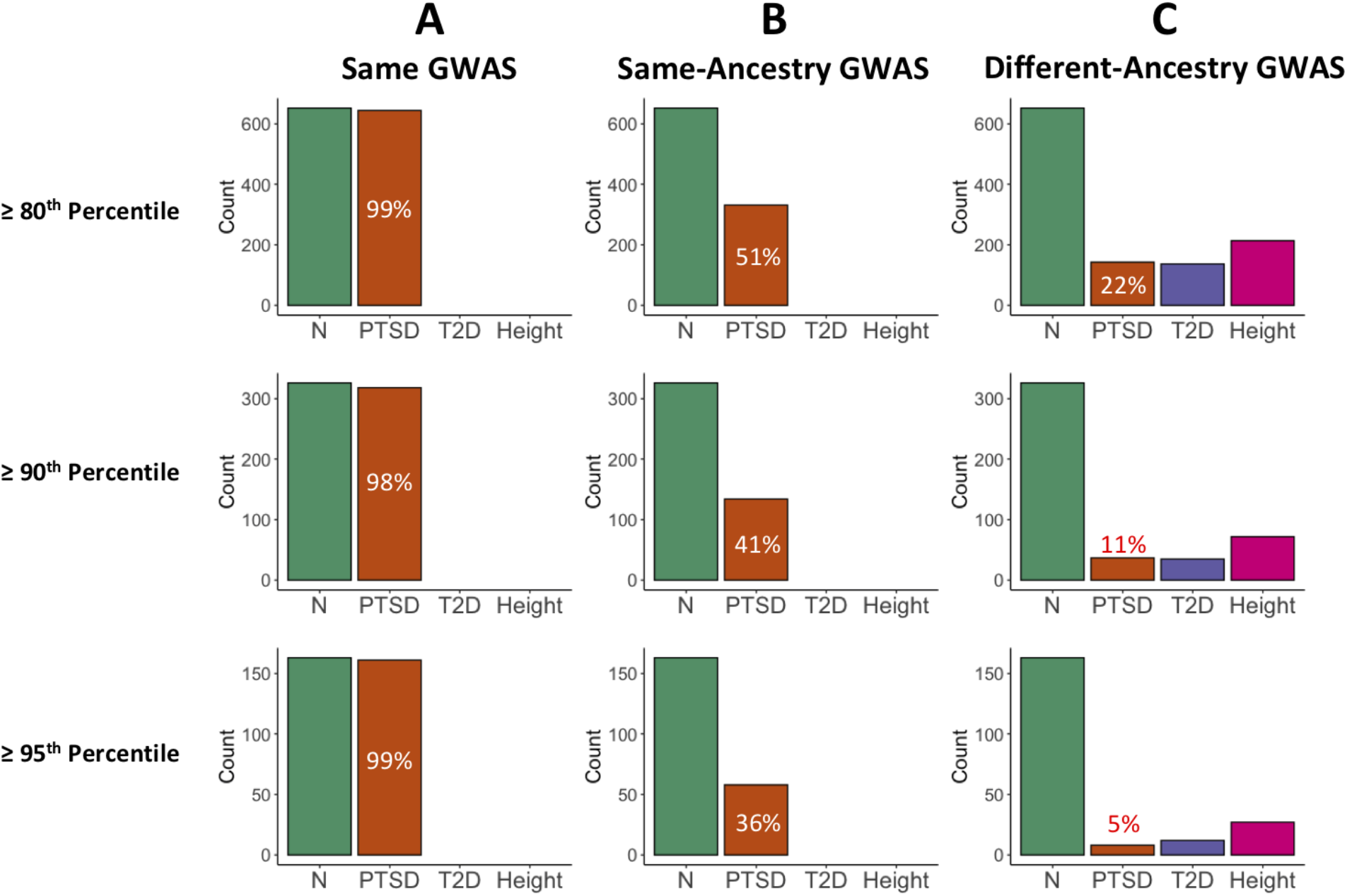
Comparison of the samples comprising the top PGS quantiles for the PNC AFR cohort. **A.** The samples located at the top 20%, 10%, and 5% of the PTSD PGS distribution were virtually the same when PGS were computed twice using the same discovery GWAS. For example, 644 out of the 652 samples (98.7%) at or above the 80^th^ percentile were the same between the two batches of PGS. **B.** The overlap between samples at all three quantiles dropped substantially when the PGS computed from the AFR PGC Freeze1 PTSD discovery GWAS^36^ were compared to those computed from the AFR Freeze2 PTSD discovery GWAS,^37^ with the degree of overlap being reduced at higher quantiles. **C.** The degree of overlap was further reduced when comparing PGS computed from an AFR-ancestry discovery GWAS to those computed from an EUR-ancestry GWAS for PTSD,^37^ T2D,^39; 40^ and height.^42^ For context, the green bars depict the number of samples included at or above the 80^th^ percentile (*n* = 652), 90^th^ percentile (*n* = 326), and 95^th^ percentile (*n* = 163). Full results can be found in Table S6.

**Figure 9.**
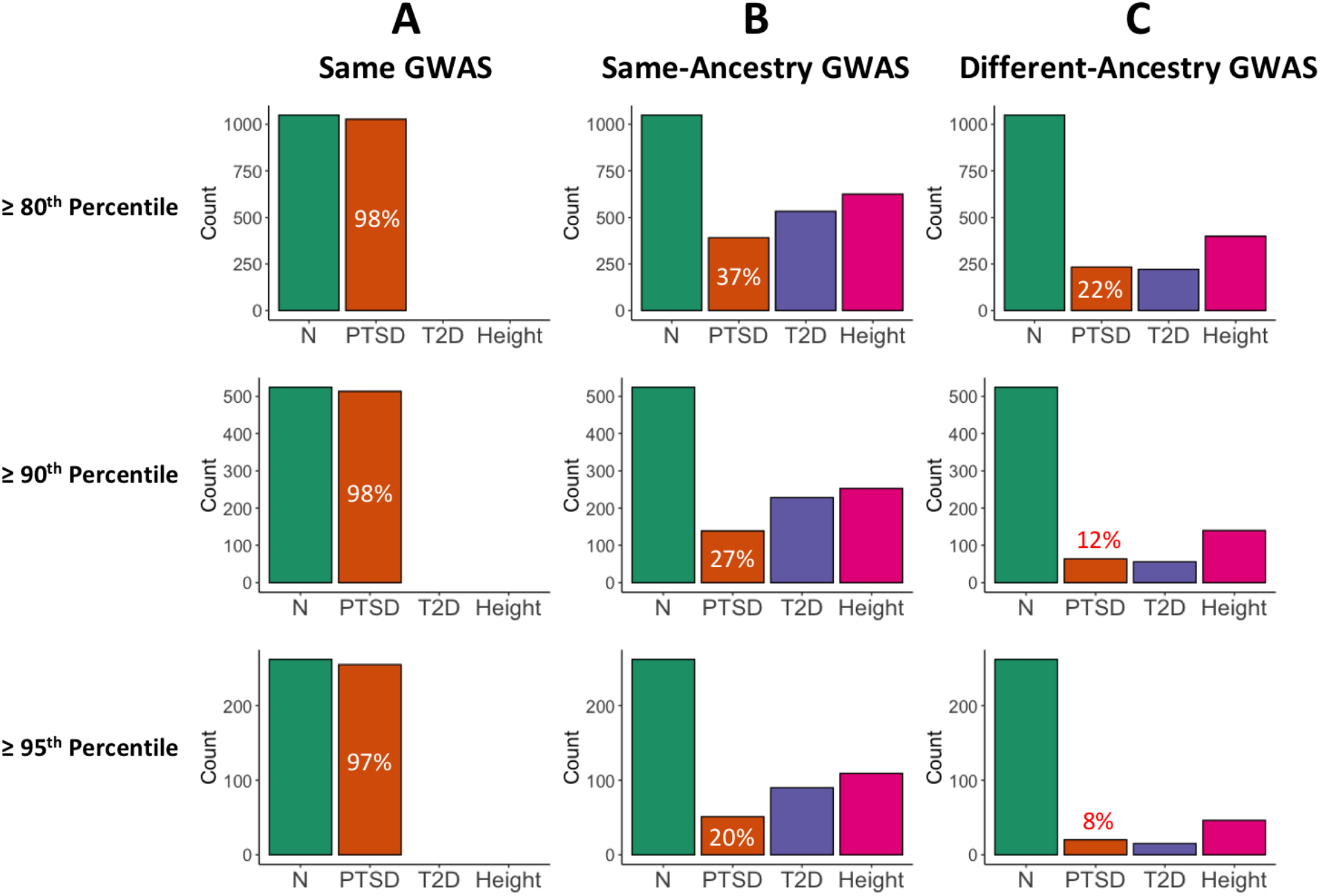
Comparison of the samples comprising the top PGS quantiles for the PNC EUR cohort. **A.** The EUR samples located within the top 20%, 10%, and 5% of the PTSD PGS distribution were nearly the same when PGS were computed twice using the same EUR discovery GWAS.^37^ For example, 1026 out of the 1048 samples (97.9%) at or above the 80^th^ percentile were the same between the two runs of PRS-CS. **B.** The overlap between samples at all three quantiles dropped substantially when the PGS computed from two different EUR discovery GWAS were compared for PTSD,^36; 37^ T2D,^38; 39^ and height.^41; 42^ **C.** The degree of overlap was dramatically reduced when comparing PGS computed from an AFR-ancestry discovery GWAS to those computed from an EUR-ancestry GWAS for PTSD,^37^ T2D,^39; 40^ and height.^41; 42^ Green bars depict the number of samples included at or above the 80^th^ percentile (*n* = 1048), 90^th^ percentile (*n* = 524), and 95^th^ percentile (*n* = 262). Full results can be found in Table S8.

The proportional overlap decreases if we consider PGS computed from two different same-ancestry discovery GWAS. For the PNC AFR-ancestry cohort (Figure 8B), adjusted standardized PGS computed from the Freeze 1^36^ and Freeze 2^37^ PTSD AFR-ancestry discovery GWAS (Figure 8B) had 53.6% of individuals in common at or above the 80^th^ percentile, 47.5% at or above the 90^th^ percentile, and 36.3% at or above the 95^th^ percentile. The decrease in proportional overlap was even more pronounced for PGS computed from two different EUR-ancestry GWAS for the PNC EUR-ancestry cohort (Figure 9B). The proportion of overlap became progressively smaller as we considered progressively higher percentiles for PTSD, T2D, and height. Moreover, the amount of overlap was greatest for height and smallest for PTSD at each of the percentiles that we considered.

Proportional overlap was even more dramatically decreased when we compared the top quantiles of the PGS that had been computed from an AFR-ancestry discovery GWAS to those that had been computed from a EUR-ancestry discovery GWAS (Figures 8C and 9C). Note that Figures 8 and 9 only include comparisons for PNC between the PGS computed using the newer discovery GWAS if there was more than one comparison possible; full results are presented in the supplement (Tables S10-S13). For the AFR cohort of PNC, the proportional overlap ranged from a low of 4.91% for different-ancestry PTSD PGS at the 95^th^ percentile to a high of 32.8% for different-ancestry height PGS at the 80^th^ percentile, whereas the proportional overlap for the PNC EUR cohort ranged from 3.82% for different-ancestry T2D PGS at the 95^th^ percentile to 38.1% for height PGS at the 80^th^ percentile. For both the EUR and AFR cohorts, the general pattern is that proportional overlap is largest for different-ancestry PGS at the 80^th^ percentile and smallest at the 95^th^ percentile. Within a given percentile, the proportional overlap is largest for height and smallest for either PTSD or T2D.

### SNP Heritability

We hypothesized that PGS would be more stable for traits with discovery GWAS that explain more of the genomic variance. As such, we used LDSC to compute the SNP heritability for each of the EUR-ancestry discovery GWAS that we used to compute PGS (Figure 10). The genetic correlation calculated by LDSC for each pair of GWAS essentially perfect, with the lowest *r_g_* = 0.9225 ± 0.1807 for PTSD. We found that SNP heritability on the observed scale was highest for height^42^ (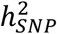 = 0.697 ± 0.067) and lowest for PTSD^37^ (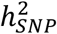 = 0.017 ± 0.003). On the liability scale (which is only relevant for the two disease traits), SNP heritability was substantially higher for T2D than for PTSD (Table S14). On both the observed and liability scales, there was a clear difference between the SNP heritability of the two discovery GWAS that were used to compute PGS for each trait. For height and T2D, the SNPs included in the newer GWAS explained more phenotypic variance than did those in the older GWAS, but the opposite was true for PTSD.

**Figure 10.**
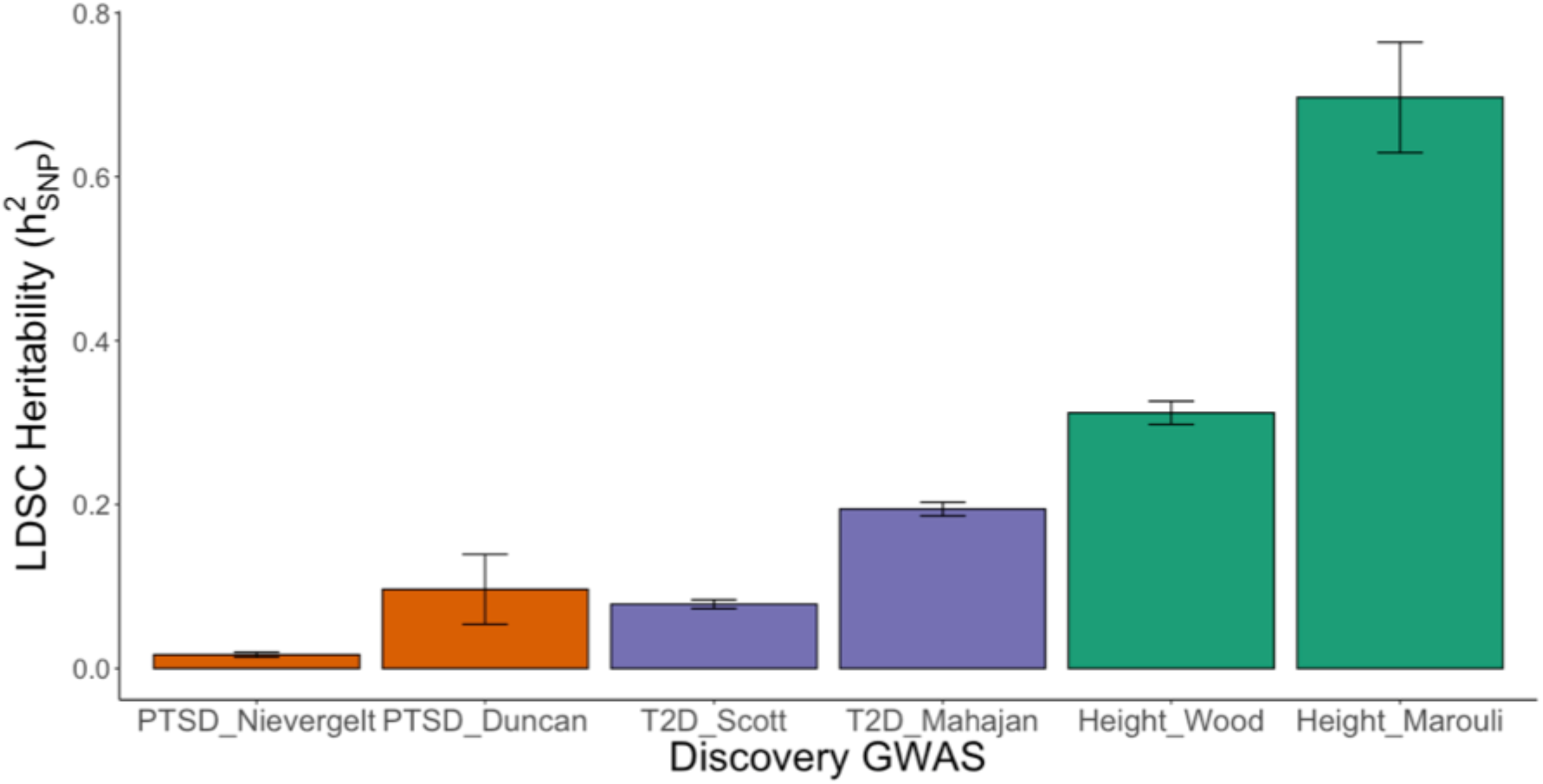
SNP heritability 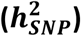 observed-scale estimates calculated using LDSC for the European-ancestry height, PTSD, and T2D discovery GWAS that were used to compute PGS. Error bars denote standard error computed via a block-jackknife procedure. Heritability was highest for height, with the newer discovery GWAS by Marouli et al. (2017)^42^ showing greater heritability than the GWAS by Wood et al. (2014)^41^. The newer T2D discovery GWAS by Mahajan et al. (2018)^39^ likewise showed greater heritability than the older GWAS by Scott et al. (2017).^38^ PTSD showed the lowest heritability, with the older discovery GWAS by Duncan et al. (2018)^36^ having greater heritability than the GWAS by Nievergelt et al. (2019).^37^

## Discussion

We have demonstrated that the PGS computed from different discovery GWAS have little to no correlation at the level of the individual patient. While the correlation is better for highly heritable anthropometric traits (e.g., height), the lack of correlation for medical and psychiatric disorders like T2D and PTSD underscores the need to proceed cautiously with integrating PGS into precision medicine applications.

This lack of correlation is especially noteworthy given that it was observed for PGS computed using meta-GWAS that were produced by the PGC,^36;37^ DIAGRAM,^38;39^ and GIANT^39;41^ consortia. The fact that even same-ancestry meta-GWAS computed by the same consortia using overlapping samples and SNPs (Table 1) would yield PGS that lack meaningful correlation at the individual level raises serious concerns. If PGS are going to be used clinically, then they need to be reproducible. Furthermore, judging PGS quality based primarily on population-level analytics, such as the area under the receiver operator characteristic curve (AUC), that depend on known case-control status will not be an effective approach for judging the validity of PGS computed for individual patients who do not necessarily have known phenotypes.^35;56^

Even if stand-alone PGS are not yet useful clinically, they could still be used to help identify those patients at highest disease risk.^57^ For instance, PGS for psychiatric traits could be used in conjunction with environmental factors to identify adolescents most at risk for developing psychosis and other mental health disorders.^16^ We are actively pursuing such applications with the PNC and ABCD cohorts and have found that ancestry-specific PTSD PGS do indeed add predictive value to models that include other non-genetic factors.^58^ Nonetheless, we caution that it is dangerous to rely solely on PGS quantiles to identify at-risk patients. Different discovery GWAS yielded PGS that did not identify the same patients at the top quantiles of the distribution, and the amount of overlap decreased as higher quantiles were considered (Figures 8 and 9; Tables S10-S13). Hence, the instinctive decision to focus only on the upper tail of the PGS distribution will not mitigate the lack of PGS stability across different discovery GWAS.

We chose to use the Bayesian PRS-CS Python package to compute PGS for this study. It has been demonstrated^4^ that Bayesian methods generally yield more predictive PGS than those produced via traditional *P*-value thresholding approaches. The advantage of PRS-CS over other Bayesian methods is that it employs a very robust Strawderman-Berger continuous shrinkage prior rather than a discrete mixture prior, which allows for more accurate multivariate modeling of local LD in the polygenic prediction.^3^ When enough MCMC iterations are used to ensure convergence of the underlying Gibbs sampler algorithm, PRS-CS yields very consistent posterior effects (Figure 2). PGS computed using the same discovery GWAS are highly correlated when computed using multiple PRS-CS runs (Figure 3), and others have previously shown that PGS computed from the same discovery GWAS are strongly correlated when computed using PRS-CS and other Bayesian and non-Bayesian approaches.^4^ Hence, the lack of PGS stability across discovery GWAS that we report here cannot be attributed to the stochastic nature of Bayesian methods; there must be differences between the discovery GWAS.

By choosing to use multiple generations of GWAS produced by the same consortia, we hoped to minimize potential methodological differences between the same-trait meta-GWAS. As expected, the genetic correlation between each pair of same-trait GWAS was nearly perfect, no doubt due to the large overlap between SNPs and samples within each pair (Table 1). Initially, we had assumed that the newer GWAS in each pair would be the “better” GWAS since we thought that the larger sample size would yield more explanatory power. We cannot rule out this possibility for the T2D and height GWAS; in both cases, the newer EUR GWAS had higher SNP heritability than the older one (Figure 10). However, given that the opposite was true for the two PTSD GWAS, we have come to believe that bigger GWAS are not necessarily better.

Phenotyping quality should not be overlooked when choosing a discovery GWAS.

Consortia have developed QC pipelines, such as RICOPILI,^59^ to harmonize genomic data from multiple cohorts prior to meta-analyzing. However, ensuring consistent phenotyping poses a continuing challenge, especially for case-control meta-GWAS analyses of non-continuous psychiatric traits like PTSD. The PGC Freeze 2 EUR-ancestry PTSD GWAS^37^ had a large increase in sample size due to the addition of samples from the UK Biobank, yet the SNP heritability went down compared to the Freeze 1 GWAS.^36^ One explanation could be that the UK Biobank PTSD phenotypes were derived from self-reported questionnaires, which are less reliable than the ascertainment procedures used by the smaller studies that comprised the original metaanalysis.^60^

It is not surprising that the two height GWAS had higher SNP heritability than the PTSD and T2D GWAS (Figure 10; Table S14). Height is an easily measured quantitative trait that is less susceptible to ascertainment bias than qualitative disease traits. Furthermore, environmental factors make substantial contributions to the development of both PTSD^61^ and T2D.^62^ Even so, LDSC gave an unusually high estimate of SNP heritability (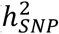 = 0.6967 ± 0.0674) for the newer height GWAS.^42^ While it is possible that the LDSC calculations could have been biased due to the small number of SNPs, we believe that a plausible explanation lies in the design of this GWAS. Specifically, the newer height GWAS included a small number of targeted rare and low-frequency SNPs (MAF between 0.1% and 4.8%) on a specially designed exome array rather than casting the same wide net as the earlier GWAS. This modification coupled with a substantially increased sample size and an easily ascertained quantitative trait could have yielded this improvement in explanatory power. It is tantalizing to speculate that a new class of genomic association studies that incorporate a smaller number of low-frequency and rare variants with large effects could improve our understanding of the genetic basis of complex traits beyond what can be inferred from large-scale GWAS alone.^63^ Such studies will be facilitated by the recent release of the Trans-Omics for Precision Medicine (TOPMed)^64^ Imputation Server, which should yield improved imputation accuracy for low-frequency and rare variants. Given that SNP heritability for psychiatric disorders generally underestimates the heritability calculated from family studies,^65^ it could be worthwhile to adopt this type of targeted approach coupled with more closely harmonized phenotyping for future genomic association studies of PTSD and other neuropsychiatric traits.

The previous heritability discussion is limited to the EUR-ancestry discovery GWAS. We chose not to use LDSC to compute SNP heritability for the AFR-ancestry GWAS due to evidence that it yields biased estimates for admixed populations.^55^ Given the relatively smaller sample sizes of the AFR GWAS (Table 1), you might expect that they explained less trait variance than the corresponding EUR GWAS. However, this was not necessarily the case. Using individual-level data, the Freeze 2 PTSD GWAS authors^37^ demonstrated that SNP heritability was comparable for EUR- and AFR-ancestry males on the liability scale, and it was higher for AFR-ancestry females than for EUR-ancestry females and both groups of males. Likewise, the relatively stronger correlation between pairs of same-ancestry AFR PTSD PGS (Figure 4) and the higher degree of overlap between individuals at upper end of the PGS distribution (Figure 8B; Tables S10 and S11) as compared to same-ancestry EUR PTSD PGS (Figures 5 and 9B; Tables S12 and S13) is consistent with the possibility that the smaller AFR-ancestry PTSD GWAS (neither of which included UK Biobank samples) explained more variance than the EUR-ancestry GWAS did. Ideally, future work will be directed towards developing methodology to calculate unbiased estimates of SNP heritability for AFR-ancestry and other admixed populations from GWAS summary statistics. Popcorn,^66^ a Python package for calculating trans-ethnic genetic correlation from GWAS summary statistics, provides a preliminary step in this direction, as does cov-LDSC.^55^

Our results add to the growing body of evidence that PGS should be computed from an ancestrally matched discovery GWAS. It is well established that EUR-ancestry GWAS typically yield PGS that are less predictive for AFR and other non-EUR ancestry groups.^19;21;25;30;31;34;67–71^ We have further demonstrated that PGS computed from same-ancestry GWAS for PTSD and T2D are uncorrelated with those computed from different-ancestry GWAS for both AFR- and EUR-ancestry study participants (Figures 6 and 7), and we also found that there is very little overlap between the individuals in the upper tails of the PGS distributions computed using EUR-ancestry GWAS as compared to those computed using AFR-ancestry GWAS (Figures 8c and 9c; Tables S10-S13). Given the dearth of AFR-ancestry and other non-EUR-ancestry discovery GWAS, our results underscore the urgent need for more high-powered GWAS analyses to be run for non-EUR ancestry populations.

We chose to study PTSD, T2D, and height because all three traits had publicly available GWAS for both EUR- and AFR-ancestry populations. Of these three, PTSD was the only trait that had two AFR-ancestry GWAS available for comparison purposes. While we focused our current work on the EUR- and AFR-ancestry individuals in the PNC and ABCD cohorts, we hope that methodology and GWAS data will soon exist to make it possible expand our analyses to the admixed American (AMR) and other ancestral groups that are also included in these cohorts (Figure 1). The recent release of PRS-CSx^72^ will make it possible to use discovery GWAS that include a combination of EAS-, AFR-, and EUR-ancestry samples. Although it offers an improvement over the current requirement that the discovery GWAS be limited to only one of these three ancestry groups, PRS-CSx still does not enable analyses of admixed samples from other genetic backgrounds.

Ultimately, we envision a future where genetic ancestry will not be a necessary consideration before computing PGS. Given that genetic ancestry is continuous, it is rather artificial to assign samples to discrete ancestry groups.^26^ Within the AFR-ancestry group alone, there is an enormous degree of genetic diversity.^31;73^ We controlled for such diversity by calculating PGS separately for each ancestry group and then regressing out within-ancestry principal components from the standardized PGS. We are optimistic that new methods that incorporate local ancestry^33;74^ will eventually allow us to embrace this diversity and compute stable, accurate PGS for admixed populations. Increasingly economical whole-genome sequencing^75^ coupled with expanded (i.e., less Eurocentric) genotyping arrays^71^ and improved imputation to diverse reference panels from TOPMed^64^ should also facilitate the further development of inclusive approaches, such as BOLT-LMM,^76;77^ trans-ethnic GWAS,^78^ and multiethnic PGS.^32^ While it certainly would be easier to continue to focus PGS development on EUR-ancestry populations, we do so at the grave risk of further exacerbating the inequities in medical care between EUR-ancestry populations and the rest of the world.^29;79^

## Description of Supplemental Data

Supplemental data include six figures and fourteen tables.

## Acknowledgments

This work was supported by the following National Institute of Mental Health grants: 5U01MH119690-02, 3U01MH119690-02, 5K23MH120437-02, and 5R01MH119219-02.

Data used in the preparation of this article were obtained from the Adolescent Brain Cognitive Development^SM^ (ABCD) Study (https://abcdstudy.org), held in the NIMH Data Archive (NDA). This is a multisite, longitudinal study designed to recruit more than 10,000 children age 9-10 and follow them over 10 years into early adulthood. The ABCD Study® is supported by the National Institutes of Health and additional federal partners under award numbers U01DA041048, U01DA050989, U01DA051016, U01DA041022, U01DA051018, U01DA051037, U01DA050987, U01DA041174, U01DA041106, U01DA041117, U01DA041028, U01DA041134, U01DA050988, U01DA051039, U01DA041156, U01DA041025, U01DA041120, U01DA051038, U01DA041148, U01DA041093, U01DA041089, U24DA041123, U24DA041147. A full list of supporters is available at https://abcdstudy.org/federal-partners.html. A listing of participating sites and a complete listing of the study investigators can be found at https://abcdstudy.org/consortium_members/. ABCD consortium investigators designed and implemented the study and/or provided data but did not necessarily participate in the analysis or writing of this report. This manuscript reflects the views of the authors and may not reflect the opinions or views of the NIH or ABCD consortium investigators.

Support for the collection of the data for Philadelphia Neurodevelopment Cohort (PNC) was provided by grant RC2MH089983 awarded to Raquel Gur and RC2MH089924 awarded to Hakon Hakonarson. Subjects were recruited and genotyped through the Center for Applied Genomics (CAG) at The Children’s Hospital in Philadelphia (CHOP). Phenotypic data collection occurred at the CAG/CHOP and at the Brain Behavior Laboratory, University of Pennsylvania.

## Web Resources

McCarthy Group imputation checking perl script

(https://www.well.ox.ac.uk/~wrayner/tools/index.html#Checking)

McCarthy Group genotyping chip strand and build files

(https://www.well.ox.ac.uk/~wrayner/strand/)

Plink 1.9

(https://www.cog-genomics.org/plink/1.9/)

Bcftools

(https://github.com/samtools/bcftools)

Michigan Imputation Server

(https://imputationserver.sph.umich.edu/index.html)

KING: Kinship-based Inference for GWAS

(http://people.virginia.edu/~wc9c/KING/index.html)

The R Project for Statistical Computing

(https://www.r-project.org)

R package ‘e1071’

(https://cran.r-project.org/web/packages/e1071/e1071.pdf)

R package ‘qqman’

(https://cran.r-project.org/web/packages/qqman/vignettes/qqman.html)

PRS-CS

(https://github.com/getian107/PRScs)

LDSC

(https://github.com/bulik/ldsc)

DIAGRAM GWAS summary statistics

(http://diagram-consortium.org/downloads.html)

GIANT GWAS summary statistics

(https://portals.broadinstitute.org/collaboration/giant/index.php/GIANT_consortium_data_files)

Psychiatric Genomics Consortium GWAS summary statistics

(https://www.med.unc.edu/pgc/data-index/)

## Data and Code Availability

The PNC and ABCD genomic datasets used in this study are available by application from dbGaP (phs00060) and NDAR (NDA #2573), respectively. All discovery GWAS summary statistics and software used in this study are publicly available; see Web Resources for access information.

