## Supplement for "Stability of Polygenic Scores Across Discovery Genome-Wide Association Studies"

**Supplemental Methods**

Pre-imputation quality control (QC) and imputation were done separately for each of the fifteen genotyping array batches that comprise the Philadelphia Neurodevelopmental Cohort (PNC) dataset. Due to the substantial variation in SNPs contained on the arrays and the numbers of samples genotyped on each array (Table S1), we imputed each array batch separately to the 1000 Genomes Mixed/Other reference panel rather than assigning ancestry prior to imputation. The fifteen batches were merged by chromosome after imputation, and post-imputation QC was run on the merged chromosome files.

Genetic ancestry was inferred by KING[1] from the principal components (PCs) derived using multi-dimensional scaling (MDS) of the hard-call dataset that was produced with PLINK 1.9 after concatenating the post-imputation-QC chromosome files. Each array batch included samples from more than one ancestry group (Table S2), thus validating our decision to not impute the array batches to specific ancestry panels.

After splitting the dataset into European-American (EUR) and African-American (AFR) cohorts, we ran a second round of unprojected MDS for each cohort separately. The first ten PCs were later regressed out of the standardized polygenic scores (PGS) to correct for both population structure and array batch effects. Batch effects, which were captured by PC2, were especially pronounced for the AFR samples that were genotyped on array_01 and array_07 (Figure S1). There were no obvious batch effects visible in the second-round PC plot for the EUR samples (Figure S2).

We further analyzed the PNC array batch effects by running a series of logistic-regression GWAS with a single array as a dummy "case" and the other arrays as dummy "controls" within the

EUR and AFR subgroups. We only used arrays that had been used to genotype at least 100 samples as "cases." For the AFR subgroup, these arrays included array_01, array_03, array_04, array_05, array_06, and array_08; the arrays for the EUR subgroup that were run as "cases" were array_03, array_04, array_05, array_06, and array_09. The GWAS were run in PLINK 1.9 both including and not including the first 10 second-round ancestry PCs as covariates so that we could confirm that including the PCs would be an adequate control for array batch effects. *P*-values were generated using Fisher's exact test. We used the R package `qqman`[2] to produce Manhattan and Q-Q plots from the Bonferroni-adjusted *p*-values.

As expected, the most dramatic GWAS results were observed for AFR array_01 (Figure S3). The logistic association model without ancestry PC covariates had many "significant" SNPs, seen as many tall peaks on the Manhattan plot (Figure S3-A) and dramatic curvature on the Q-Q plot (Figure S3-B). When the first 10 PCs were included as covariates, the Q-Q plot had no deviations from the straight line (Figure S3-D). The GWAS results from the other arrays similarly yielded linear Q-Q plots when 10 within-ancestry PCs were included as covariates. Taken together, these results indicated that regressing out the first 10 within-ancestry PCs from our polygenic scores (PGS) would adequately control for any array batch effects.

The ABCD dataset was genotyped exclusively on the Affymetrix NIDA SmokeScreen array. As such, QC and imputation were done on a single dataset. Table S3 shows the ABCD SNP and sample counts before and after the pre-imputation QC. Within-ancestry PCs were computed for the AFR (Figure S3) and EUR (Figure S4) subsets of the ABCD dataset.

PRS-CS[3] requires a single SNP sample size as an input. Given that most of our discovery GWAS were meta-GWAS that were comprised of individual studies that varied in terms of their sample size and the SNPs they included, the effective sample size often varied considerably between SNPs. To account for this reality, we examined the distribution of SNP sample sizes in R and excluded SNPs that had sample sizes that were less than half of the maximum SNP sample size. Of the remaining SNPs, the median SNP sample size was used as the PRS-CS sample size input.

As an example, consider the Freeze 2 EUR PTSD GWAS produced by the Psychiatric Genomics Consortium (PGC).[4] This meta-GWAS includes 9,766,174 SNPs with effective sample sizes that range from 17,559.4 to 70,237.5 (Figure S6-A). Given that PRS-CS uses only those SNPs that overlap with both the relevant LD panel and the dataset, we started by retaining only the 1,116,862 SNPs that were present in the EUR LD panel. These SNPs also had effective sample sizes ranging from 17,559.4 to 70,237.5 (Figure S6-B). After we removed SNPs with effective sample sizes that were less than 35,000, the remaining 1,113,044 SNPs had effective sample sizes that ranged from 38,250.5 to 70,237.5 (Figure S6-C), with a median of 70,237.5. This median value was truncated to 70,237 and used as the SNP sample size when we ran PRS-CS. We made similar sample size determinations for the other discovery GWAS.

**Supplemental Data**

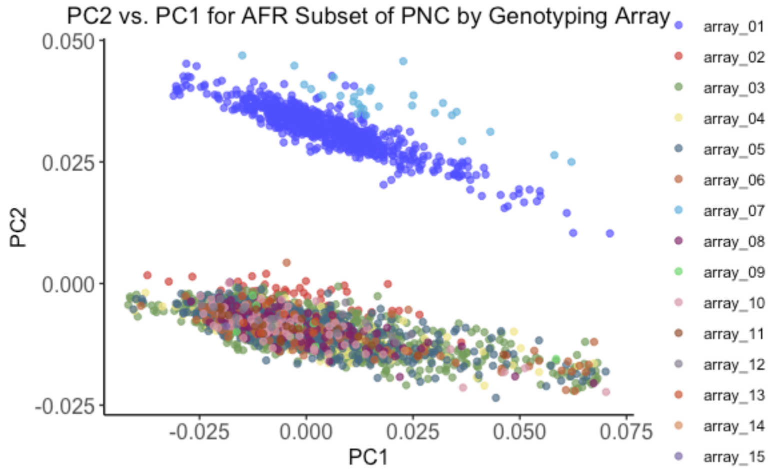


**Figure S1.** Within-ancestry PC2 vs. PC1 for the AFR subset of PNC with samples color-coded by their genotyping array batch. PC2 captures an array batch effect that is most pronounced for array_01 and array_07.

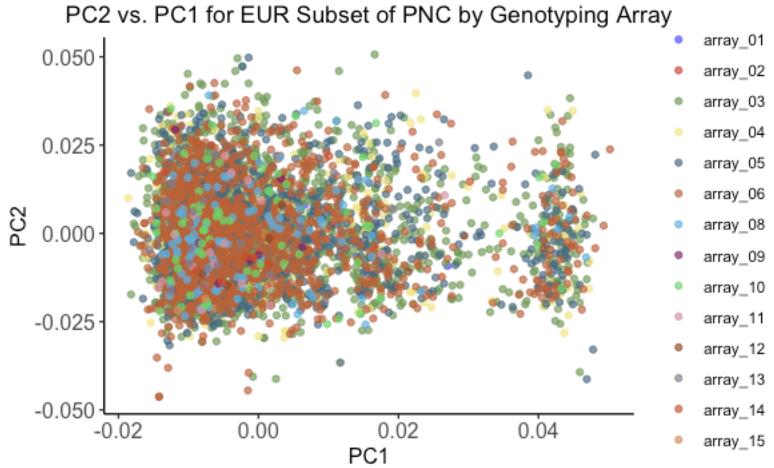


**Figure S2.** Within-ancestry PC2 vs. PC1 for the EUR subset of PNC with samples color-coded by their genotyping array batch.

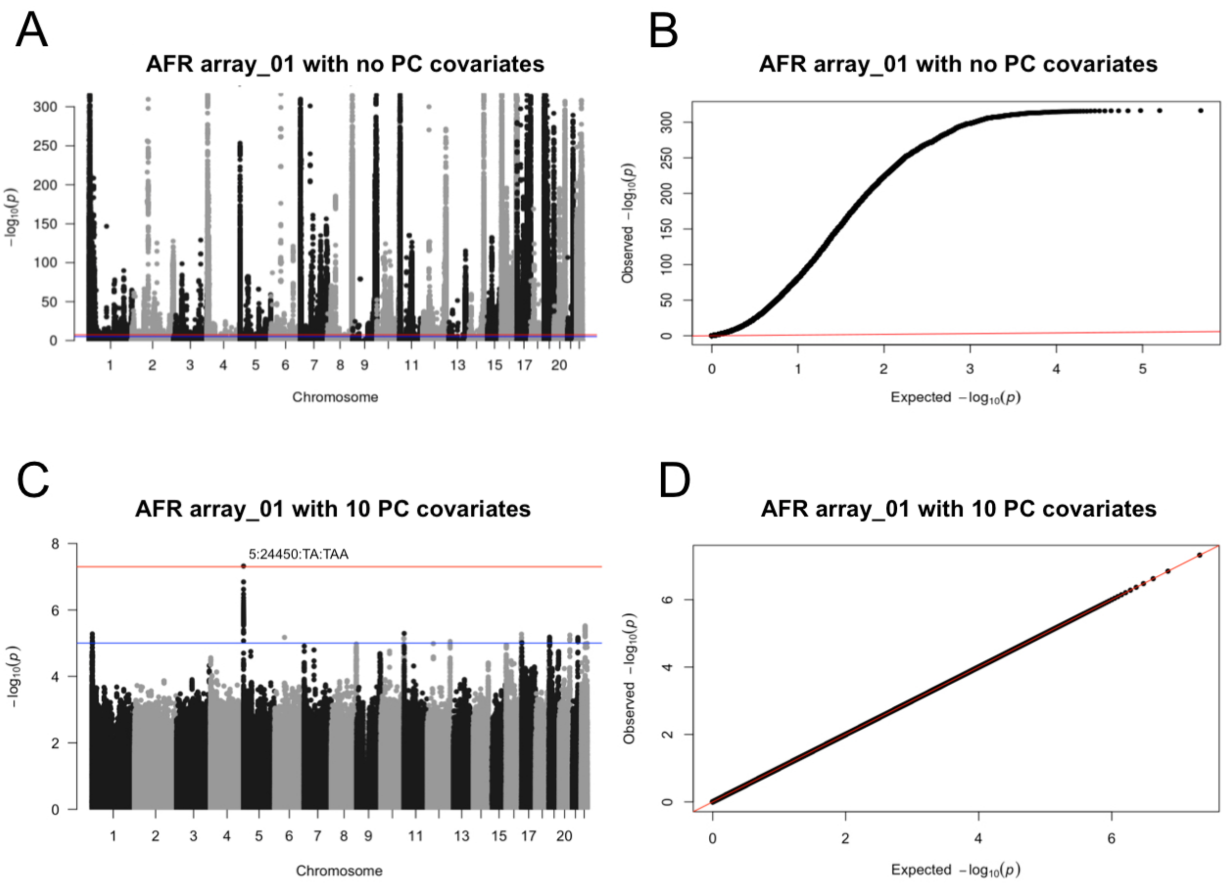


**Figure S3.** Illustration of PNC batch effects for AFR array_01. (A) The Manhattan plot showed many highly significant SNPs when the ancestry PCs were not included as covariates. (B) When PCs were not included as covariates, the Q-Q plot deviated substantially from the expected straight line. (C) When 10 PCs were included as covariates, no significant peaks remained in the Manhattan plot. (D) With 10 PCs included as covariates, the Q-Q plot of observed $-\log_{10}(p)$ versus expected $-\log_{10}(p)$ followed a straight line.

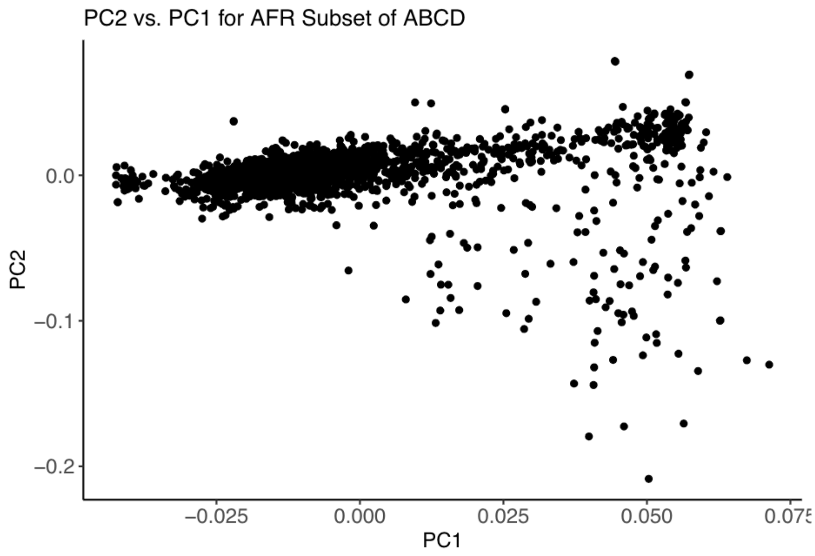


**Figure S4.** Within-ancestry PC2 vs. PC1 for the AFR subset of the ABCD dataset.

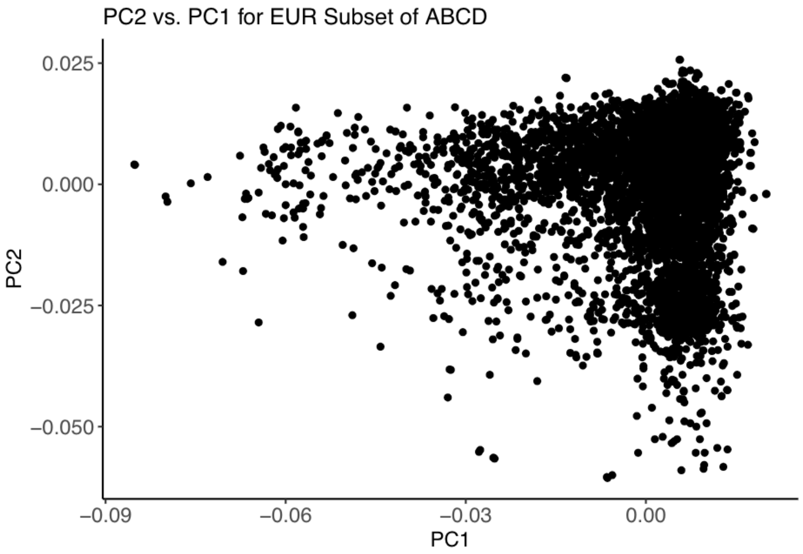


**Figure S5.** Within-ancestry PC2 vs. PC1 for the EUR subset of the ABCD dataset.

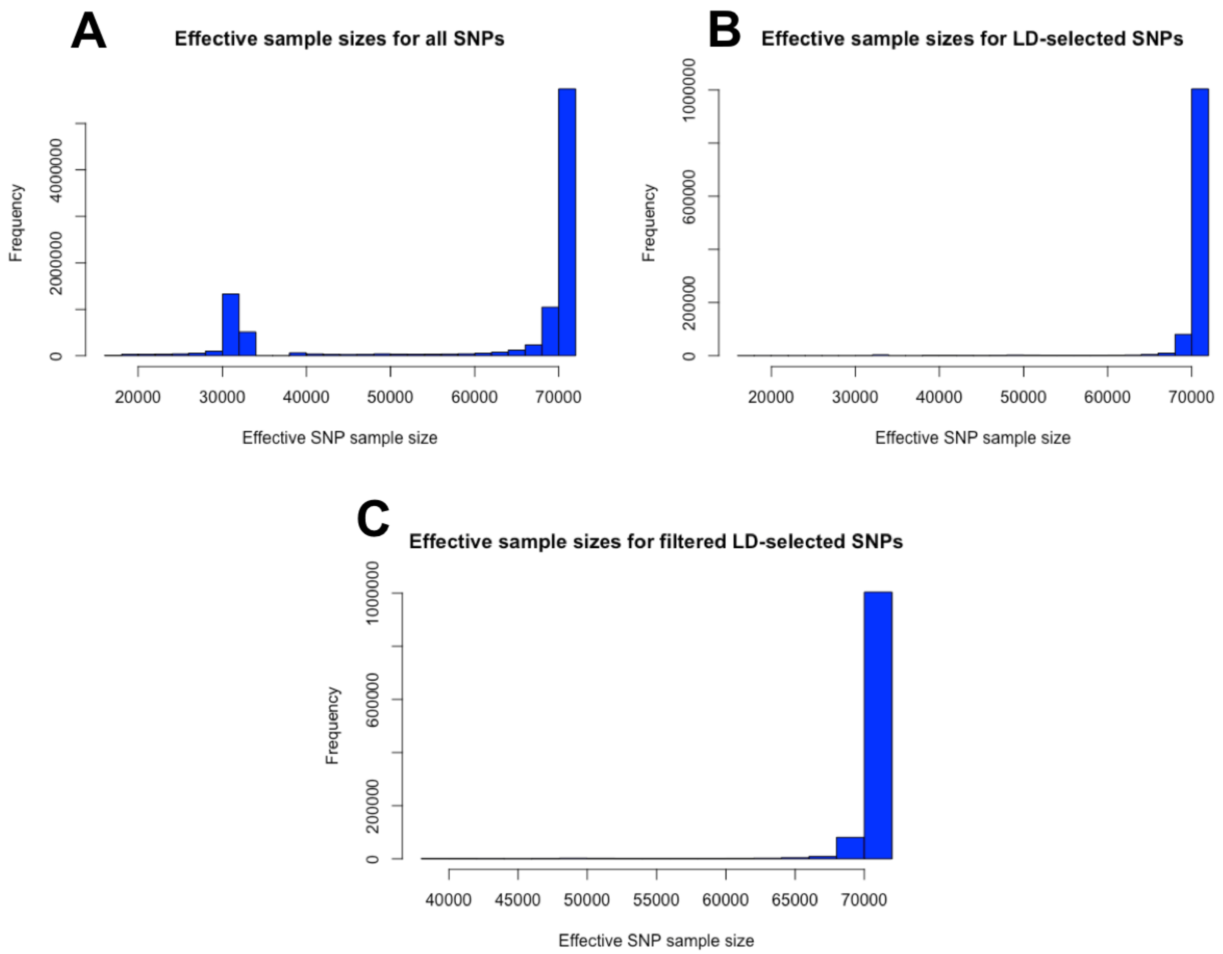


**Figure S6.** SNP sample size determination for the PTSD Freeze 2 EUR meta-GWAS[4]. (A) This meta-GWAS contained 9,766,174 SNPs with effective sample sizes that ranged from 17,559.4 to 70,237.5. (B) The 1,116,862 SNPs that were present in the PRS-CS EUR LD panel had this same range of effective sample sizes. (C) Filtering to retain only LD-selected SNPs with effective samples sizes of at least 35,000 resulted in 1,113,044 SNPs with effective sample sizes between 38,250.5 and 70,237.5. The median effective SNP sample size of 70,237.5 for these filtered SNPs was truncated to 70,237 and used as the SNP sample size in PRS-CS.

**Table S1. PNC sample and SNP counts by genotyping array before and after pre-imputation QC.**

| Array Code | dbGaP Filename | No. of Samples Pre-QC | No. of Samples Post-QC | No. of SNPs Pre-QC | No. of SNPs Post-QC |
|---|---|---|---|---|---|
| array_01 | phg000381.v2.NIMH_NeurodevelopmentalGenomics.genotype-calls-matrixfmt.Axiom.c1.GRU-NPU/GO_Axiom | 722 | 719 | 567,096 | 472,217 |
| array_02 | phg000381.v2.NIMH_NeurodevelopmentalGenomics.genotype-calls-matrixfmt.Genome-Wide_Human_SNP_Array_6.0.c1.GRU-NPU/GO_Affy60 | 66 | 66 | 909,622 | 725,897 |
| array_03 | phg000381.v2.NIMH_NeurodevelopmentalGenomics.genotype-calls-matrixfmt.Human610-Quadv1_B.c1.GRU-NPU/GO_Quad_5removed | 3,802 | 3,789 | 620,901 | 573,487 |
| array_04 | phg000381.v2.NIMH_NeurodevelopmentalGenomics.genotype-calls-matrixfmt.HumanHap550_v1.c1.GRU-NPU/GO_v1_1removed | 555 | 552 | 555,352 | 533,783 |
| array_05 | phg000381.v2.NIMH_NeurodevelopmentalGenomics.genotype-calls-matrixfmt.HumanHap550_v3.c1.GRU-NPU/GO_v3_1removed | 1,913 | 1,893 | 561,466 | 541,643 |
| array_06 | phg000381.v2.NIMH_NeurodevelopmentalGenomics.genotype-calls-matrixfmt.HumanOmniExpress.c1.GRU-NPU/GO_Omni | 1,657 | 1,654 | 733,202 | 693,213 |
| array_07 | phg000661.v1.NIMH_NeurodevelopmentalGenomics_v2.genotype-calls-matrixfmt.Axiom.c1.GRU-NPU/GO_Axiom_set2 | 40 | 32 | 567,096 | 298,921 |
| array_08 | phg000661.v1.NIMH_NeurodevelopmentalGenomics_v2.genotype-calls-matrixfmt.Axiom.c1.GRU-NPU/GO_AxiomTx | 225 | 218 | 767,203 | 616,881 |
| array_09 | phg000661.v1.NIMH_NeurodevelopmentalGenomics_v2.genotype-calls-matrixfmt.BDCHP-1X10-HUMANHAP550.c1.GRU-NPU/GO_v1set2 | 17 | 17 | 555,352 | 490,740 |
| array_10 | phg000661.v1.NIMH_NeurodevelopmentalGenomics_v2.genotype-calls-matrixfmt.Human1M-Duov3_B.c1.GRU-NPU/GO_1MDuo | 141 | 141 | 1,199,187 | 1,040,603 |
| array_11 | phg000661.v1.NIMH_NeurodevelopmentalGenomics_v2.genotype-calls-matrixfmt.Human610-Quadv1_B.c1.GRU-NPU/GO_Quadset2 | 40 | 40 | 620,901 | 564,518 |
| array_12 | phg000661.v1.NIMH_NeurodevelopmentalGenomics_v2.genotype-calls-matrixfmt.HumanHap550_v3.c1.GRU-NPU/GO_v3set2 | 31 | 31 | 561,466 | 516,726 |
| array_13 | phg000661.v1.NIMH_NeurodevelopmentalGenomics_v2.genotype-calls-matrixfmt.HumanOmniExpress-12v1_A.c1.GRU-NPU/GO_Omniset2 | 37 | 35 | 733,202 | 674,803 |
| array_14 | phg000661.v1.NIMH_NeurodevelopmentalGenomics_v2.genotype-calls-matrixfmt.HumanOmniExpress-12v1_B.c1.GRU-NPU/GO_OMNI12v11 | 18 | 18 | 719,665 | 578,397 |
| array_15 | phg000661.v1.NIMH_NeurodevelopmentalGenomics_v2.genotype-calls-matrixfmt.HumanOmniExpressExome-8v1_A.c1.GRU-NPU/GO_OEE | 3 | 3 | 951,117 | 400,300 |

Table S2. PNC ancestry by genotyping array.

| Array | AFR | EUR | Other | Total |
|---|---|---|---|---|
| array_01 | 693 | 8 | 18 | 719 |
| array_02 | 63 | 2 | 1 | 66 |
| array_03 | 1,341 | 2,157 | 291 | 3,789 |
| array_04 | 177 | 341 | 34 | 552 |
| array_05 | 623 | 1,137 | 133 | 1,893 |
| array_06 | 112 | 1,354 | 188 | 1,654 |
| array_07 | 29 | 0 | 3 | 32 |
| array_08 | 101 | 102 | 15 | 218 |
| array_09 | 9 | 7 | 1 | 17 |
| array_10 | 62 | 69 | 10 | 141 |
| array_11 | 20 | 18 | 2 | 40 |
| array_12 | 19 | 8 | 2 | 29 |
| array_13 | 9 | 18 | 8 | 35 |
| array_14 | 1 | 16 | 1 | 18 |
| array_15 | 1 | 2 | 0 | 3 |
| Total | **3,260** | **5,239** | **707** | **9,206** |

Table S3. ABCD sample and SNP counts before and after Pre-Imputation QC.

| Number of Samples Pre-QC | Number of Samples Post-QC | Number of SNPs Pre-QC | Number of SNPs Post-QC |
|---|---|---|---|
| 10,461 | 10,318 | 517,724 | 483,017 |

**Table S4. PTSD PGS correlations for PNC EUR cohort when limited to one sample per family versus including all samples.**

| Comparison | One sample per family $n$ = 4928 | All EUR samples $n$ = 5239 |
|---|---|---|
| PRS-CS replication using same discovery GWAS (PGC Freeze 2)[4] | $r$ = 0.9994 ($t$ = 2007 , $P$ < 2e-16) | $r$ = 0.9994 ($t$ = 2057, $P$ < 2e-16) |
| Different discovery GWAS, same ancestry (PGC Freezes 1 and 2)[4; 5] | $r$ = 0.388 ($t$ = 29.55, $P$ < 2e-16) | $r$ = 0.392 ($t$ = 30.86, $P$ < 2e-16) |
| Different discovery GWAS, different ancestry (PGC Freeze 1)[5] | $r$ = -0.00265 ($t$ = -0.186, $P$ = 0.852) | $r$ = 0.00136 ($t$ = 0.098, P = 0.922) |
| Different discovery GWAS, different ancestry (PGC Freeze 2)[4] | $r$ = 0.0341 ($t$ =2.391, $P$ = 0.0168) | $r$ = 0.0379 ($t$ = 2.746, $P$ = 0.00605) |

PNC, Philadelphia Neurodevelopmental Cohort; PTSD, post-traumatic stress disorder; PGS, polygenic score; EUR, European-American ancestry; GWAS, genome-wide association study; PGC; Psychiatric Genomics Consortium; *r*, Pearson correlation coefficient; *t*, linear association *t*-test statistic; *P*, two-tailed *P*-value on 4926 (or 5237) degrees of freedom.
Superscripts are the reference numbers for the discovery GWAS used to calculate PGS:
[4]Nievergelt et al. (2019), [5]Duncan et al. (2018).

**Table S5. PTSD PGS correlations for PNC AFR cohort when limited to one sample per family versus including all samples.**

| Comparison | One sample per family $n$ = 2954 | All AFR samples $n$ = 3260 |
|---|---|---|
| PRS-CS replication using same discovery GWAS (PGC Freeze 2)[4] | $r$ =0.9997 ($t$ = 2055, $P$ < 2e-16) | $r$ = 0.9997 ($t$ = 2162, $P$ < 2e-16) |
| Different discovery GWAS, same ancestry (PGC Freezes 1 and 2)[4; 5] | $r$ = 0.636 ($t$ = 44.76, $P$ < 2e-16) | $r$ = 0.696 ($t$ = 55.26, $P$ < 2e-16) |
| Different discovery GWAS, different ancestry (PGC Freeze 1)[5] | $r$ = 0.0399 ($t$ = 2.171, $P$ = 0.03) | $r$ = 0.0417 ($t$ = 2.379, $P$ = 0.0174) |
| Different discovery GWAS, different ancestry (PGC Freeze 2)[4] | $r$ = 0.000732 ($t$ = 0.04, $P$ = 0.968) | $r$ = 0.00356 ($t$ = 0.203, $P$ = 0.839) |

PNC, Philadelphia Neurodevelopmental Cohort; PTSD, post-traumatic stress disorder; PGS, polygenic score; AFR, African-American ancestry; GWAS, genome-wide association study; PGC; Psychiatric Genomics Consortium; *r*, Pearson correlation coefficient; linear association *t*-test statistic, *P*, two-tailed *P*-value on 2952 (or 3258) degrees of freedom.
Superscripts are the reference numbers for the discovery GWAS used to calculate PGS:
[4]Nievergelt et al. (2019), [5]Duncan et al. (2018).

**Table S6. PGS correlations for PNC AFR cohort (*n* = 3260).**

| Comparison | PTSD | T2D | Height |
|---|---|---|---|
| PRS-CS replication using same discovery GWAS | *r* = 0.9997[4] (*t* = 2162, *P* < 2e-16) | NA | NA |
| Different discovery GWAS, same ancestry | *r* = 0.696[4; 5] (*t* = 55.26, *P* < 2e-16) | NA | NA |
| Different discovery GWAS, different ancestry | *r* = 0.0417[5] (*t* = 2.379, *P* = 0.0174) | *r* = 0.0185[6; 7] (*t* = 1.055, *P* = 0.292) | *r* = 0.287[8] (*t* = 17.09, *P* < 2e-16) |
| | *r* = 0.00356[4] (*t* = 0.203, *P* = 0.839) | *r* = 0.0432[7; 9] (*t* = 2.469, *P* = 0.0136) | *r* = 0.258[8; 10] (*t* = 15.22, *P* < 2e-16) |

PNC, Philadelphia Neurodevelopmental Cohort; PGS, polygenic score; AFR, African-American ancestry; GWAS, genome-wide association study; PTSD, post-traumatic stress disorder; T2D, type 2 diabetes; *r*, Pearson correlation coefficient; *t*, linear association *t*-test statistic; *P*, two-tailed *P*-value on 3258 degrees of freedom, NA, not applicable (analysis not run).

Superscripts are the reference numbers for the discovery GWAS used to calculate PGS:
[4]Nievergelt et al. (2019), [5]Duncan et al. (2018), [6]Mahajan et al. (2018), [7]Chen et al. (2019), [8]Marouli et al. (2017), [9]Scott et al. (2017), [10]Wood et al. (2014).

**Table S7. PGS correlations for ABCD AFR cohort (*n* = 1741).**

| Comparison | PTSD | T2D | Height |
|---|---|---|---|
| Different discovery GWAS, same ancestry | *r* = 0.657[4; 5] (*t* = 36.34, *P* < 2e-16) | NA | NA |
| Different discovery GWAS, different ancestry | *r* = -0.00320[5] (*t* = -0.133, *P* = 0.894) | *r* = 0.0219[6; 7] (*t* = 0.912, *P* = 0.362) | *r* = 0.306[8] (*t* = 13.42, *P* < 2e-16) |
| | *r* = 0.00283[4] (*t* = 0.118, *P* = 0.906) | *r* = -0.0458[7; 9] (*t* = -1.911, *P* = 0.0562) | *r* = 0.312[8; 10] (*t* = 13.68, *P* < 2e-16) |

ABCD, Adolescent Brain and Cognitive Development Study; PGS, polygenic score; AFR, African-American ancestry; GWAS, genome-wide association study; PTSD, post-traumatic stress disorder; T2D, type 2 diabetes; *r*, Pearson correlation coefficient; *t*, linear association *t*-test statistic; *P*, two-tailed *P*-value on 1739 degrees of freedom, NA, not applicable (analysis not run).

Superscripts are the reference numbers for the discovery GWAS used to calculate PGS:
[4]Nievergelt et al. (2019), [5]Duncan et al. (2018), [6]Mahajan et al. (2018), [7]Chen et al. (2019), [8]Marouli et al. (2017), [9]Scott et al. (2017), [10]Wood et al. (2014).

**Table S8. PGS correlations for PNC EUR cohort (*n* = 5239).**

| Comparison | PTSD | T2D | Height |
|---|---|---|---|
| PRS-CS replication using same discovery GWAS | *r* = 0.9994[4] (*t* = 2057, *P* < 2e-16) | NA | NA |
| Different discovery GWAS, same ancestry | *r* = 0.392[4; 5] (*t* = 30.86, *P* < 2e-16) | *r* = 0.602[6; 9] (*t* = 54.54, *P* < 2e-16) | *r* = 0.736[8; 10] (*t* = 78.78, P < 2e-16) |
| Different discovery GWAS, different ancestry | *r* = 0.00136[5] (*t* = 0.098, *P* = 0.922) | *r* = 0.0240[6; 7] (*t* = 1.739, *P* = 0.082) | *r* = 0.403[8] (*t* = 31.82, *P* < 2e-16) |
| | *r* = 0.0379[4] (*t* = 2.746, *P* = 0.00605) | *r* = 0.00528[7; 9] (*t* = 0.382, *P* = 0.703) | *r* = 0.335[8; 10] (*t* = 25.25, *P* < 2e-16) |

PNC, Philadelphia Neurodevelopmental Cohort; PGS, polygenic score; EUR, European-American ancestry; GWAS, genome-wide association study; PTSD, post-traumatic stress disorder; T2D, type 2 diabetes; *r*, Pearson correlation coefficient; *t*, linear association *t*-test statistic; *P*, two-tailed *P*-value on 5237 degrees of freedom; NA, not applicable (analysis not run).

Superscripts are the reference numbers for the discovery GWAS used to calculate PGS:
[4]Nievergelt et al. (2019), [5]Duncan et al. (2018), [6]Mahajan et al. (2018), [7]Chen et al. (2019), [8]Marouli et al. (2017), [9]Scott et al. (2017), [10]Wood et al. (2014).

**Table S9. PGS correlations for ABCD EUR cohort (*n* = 5815).**

| Comparison | PTSD | T2D | Height |
|---|---|---|---|
| Different discovery GWAS, same ancestry | *r* = 0.378[4; 5] (*t* = 31.14, *P* < 2e-16) | *r* = 0.597[6; 9] (*t* = 56.79, *P* < 2e-16) | *r* = 0.734[8; 10] (*t* = 82.46, P < 2e-16) |
| Different discovery GWAS, different ancestry | *r* = -0.00109[5] (*t* = -0.083, *P* = 0.934) | *r* = 0.0224[6; 7] (*t* = 1.71, *P* = 0.0872) | *r* = 0.404[8] (*t* = 33.69, *P* < 2e-16) |
| | *r* = 0.000867[4] (*t* = 0.066, *P* = 0.947) | *r* = 0.0188[7; 9] (*t* = 1.431, *P* = 0.152) | *r* = 0.327[8; 10] (*t* = 26.39, *P* < 2e-16) |

ABCD, Adolescent Brain and Cognitive Development Study; PGS, polygenic score; EUR, European-American ancestry; GWAS, genome-wide association study; PTSD, post-traumatic stress disorder; T2D, type 2 diabetes; *r*, Pearson correlation coefficient; *t*, linear association *t*-test statistic; *P*, two-tailed *P*-value on 5813 degrees of freedom.

Superscripts are the reference numbers for the discovery GWAS used to calculate PGS:
[4]Nievergelt et al. (2019), [5]Duncan et al. (2018), [6]Mahajan et al. (2018), [7]Chen et al. (2019), [8]Marouli et al. (2017), [9]Scott et al. (2017), [10]Wood et al. (2014).

**Table S10. Proportional overlap for PNC AFR polygenic scores (*n* = 3260).**

| **Comparison** | Top Quintile<br>(≥ 80th Percentile)<br>*n* = 652 | Top Decile<br>(≥ 90th Percentile)<br>*n* = 326 | Top Ventile<br>(≥ 95th Percentile)<br>*n* = 163 |
|---|---|---|---|
| PRS-CS replication using same discovery GWAS | PTSD:[4] 644/652 (0.987) | PTSD:[4] 318/326 (0.975) | PTSD:[4] 161/163 (0.988) |
| Different discovery GWAS, same ancestry | PTSD:[4; 5] 331/652 (0.508) | PTSD:[4; 5] 134/326 (0.411) | PTSD:[4; 5] 58/163 (0.356) |
| Different discovery GWAS, different ancestry | PTSD:[4] 143/652 (0.219)<br>PTSD:[5] 138/652 (0.212)<br>T2D:[6; 7] 137/652 (0.210)<br>T2D:[7; 9] 144/652 (0.221)<br>height:[8] 214/652 (0.328)<br>height:[8; 10] 209/652 (0.321) | PTSD:[4] 37/326 (0.113)<br>PTSD:[5] 36/326 (0.110)<br>T2D:[6; 7] 35/326 (0.107)<br>T2D:[7; 9] 41/326 (0.126)<br>height:[8] 77/326 (0.236)<br>height:[8; 10] 72/326 (0.221) | PTSD:[4] 8/163 (0.0491)<br>PTSD:[5] 12/163 (0.0736)<br>T2D:[6; 7] 12/163 (0.0736)<br>T2D:[7; 9] 13/163 (0.0798)<br>height:[8] 27/163 (0.166)<br>height:[8; 10] 31/163 (0.190) |

PNC, Philadelphia Neurodevelopmental Cohort; AFR, African-American ancestry; *n*, number of subjects; PTSD, post-traumatic stress disorder; T2D, type 2 diabetes.
Superscripts are the reference numbers for the discovery GWAS used to calculate PGS:
[4]Nievergelt et al. (2019), [5]Duncan et al. (2018), [6]Mahajan et al. (2018), [7]Chen et al. (2019), [8]Marouli et al. (2017), [9]Scott et al. (2017), [10]Wood et al. (2014).

**Table S11. Proportional overlap for ABCD AFR polygenic scores (*n* = 1741)**

| **Comparison** | Top Quintile<br>(≥ 80th Percentile)<br>*n* = 349 | Top Decile<br>(≥ 90th Percentile)<br>*n* = 175 | Top Ventile<br>(≥ 95th Percentile)<br>*n* = 88 |
|---|---|---|---|
| Different discovery GWAS, same ancestry | PTSD:[4; 5] 187/349 (0.536) | PTSD:[4; 5] 83/175 (0.475) | PTSD:[4; 5] 32/88 (0.363) |
| Different discovery GWAS, different ancestry | PTSD:[4] 66/349 (0.189)<br>PTSD:[5] 62/349 (0.178)<br>T2D:[6; 7] 76/349 (0.218)<br>T2D:[7; 9] 69/349 (0.198)<br>height:[8] 115/349 (0.330)<br>height:[8; 10] 119/349 (0.341) | PTSD:[4] 25/175 (0.143)<br>PTSD:[5] 18/175 (0.103)<br>T2D:[6; 7] 19/175 (0.109)<br>T2D:[7; 9] 19/175 (0.109)<br>height:[8] 45/175 (0.257)<br>height:[8; 10] 34/175 (0.194) | PTSD:[4] 7/88 (0.0795)<br>PTSD:[5] 4/88 (0.0455)<br>T2D:[6; 7] 10/88 (0.114)<br>T2D:[7; 9] 6/88 (0.0343)<br>height:[8] 13/88 (0.148)<br>height:[8; 10] 15/88 (0.170) |

ABCD, Adolescent Brain and Cognitive Development Study; AFR, African-American ancestry; *n*, number of subjects; PTSD, post-traumatic stress disorder; T2D, type 2 diabetes.
Superscripts are the reference numbers for the discovery GWAS used to calculate PGS:
[4]Nievergelt et al. (2019), [5]Duncan et al. (2018), [6]Mahajan et al. (2018), [7]Chen et al. (2019), [8]Marouli et al. (2017), [9]Scott et al. (2017), [10]Wood et al. (2014).

**Table S12. Proportional overlap for PNC EUR polygenic scores (*n* = 5239)**

| Comparison | Top Quintile<br>(≥ 80th Percentile)<br>*n* = 1048 | Top Decile<br>(≥ 90th Percentile)<br>*n* = 524 | Top Ventile<br>(≥ 95th Percentile)<br>*n* = 262 |
|---|---|---|---|
| PRS-CS replication using same discovery GWAS | PTSD:[4] 1026/1048 (0.979) | PTSD:[4] 513/524 (0.979) | PTSD:[4] 255/262 (0.973) |
| Different discovery GWAS, same ancestry | PTSD:[4; 5] 391/1048 (0.373)<br>T2D:[6; 9] 532/1048 (0.508)<br>height:[8; 10] 625/1048 (0.596) | PTSD:[4; 5] 139/524 (0.265)<br>T2D:[6; 9] 228/524 (0.435)<br>height:[8; 10] 253/524 (0.483) | PTSD:[4; 5] 51/262 (0.195)<br>T2D:[6; 9] 90/262 (0.344)<br>height:[8; 10] 109/262 (0.416) |
| Different discovery GWAS, different ancestry | PTSD:[4] 233/1048 (0.222)<br>PTSD:[5] 209/1048 (0.199)<br>T2D:[6; 7] 221/1048 (0.211)<br>T2D:[7; 9] 204/1048 (0.195)<br>height:[8] 399/1048 (0.381)<br>height:[8; 10] 381/1048 (0.364) | PTSD:[4] 64/524 (0.122)<br>PTSD:[5] 47/524 (0.0897)<br>T2D:[6; 7] 56/524 (0.107)<br>T2D:[7; 9] 50/524 (0.0954)<br>height:[8] 140/524 (0.267)<br>height:[8; 10] 119/524 (0.227) | PTSD:[4] 20/262 (0.0763)<br>PTSD:[5] 14/262 (0.0534)<br>T2D:[6; 7] 15/262 (0.0573)<br>T2D:[7; 9] 10/262 (0.0382)<br>height:[8] 46/262 (0.176)<br>height:[8; 10] 41/262 (0.156) |

PNC, Philadelphia Neurodevelopmental Cohort; EUR, European-American ancestry; *n*, number of subjects; PTSD, post-traumatic stress disorder; T2D, type 2 diabetes.
Superscripts are the reference numbers for the discovery GWAS used to calculate PGS:
[4]Nievergelt et al. (2019), [5]Duncan et al. (2018), [6]Mahajan et al. (2018), [7]Chen et al. (2019), [8]Marouli et al. (2017), [9]Scott et al. (2017), [10]Wood et al. (2014).

**Table S13. Proportional overlap for ABCD EUR polygenic scores (*n* = 5815)**

| Comparison | Top Quintile<br>(≥ 80th Percentile)<br>*n* = 1163 | Top Decile<br>(≥ 90th Percentile)<br>*n* = 582 | Top Ventile<br>(≥ 95th Percentile)<br>*n* = 291 |
|---|---|---|---|
| Different discovery GWAS, same ancestry | PTSD:[4; 5] 444/1163 (0.382)<br>T2D:[8; 9] 586/1163 (0.504)<br>height:[8; 10] 674/1163 (0.580) | PTSD:[4; 5] 153/582 (0.263)<br>T2D:[8; 9] 230/582 (0.395)<br>height:[8; 10] 301/582 (0.517) | PTSD:[4; 5] 67/291 (0.230)<br>T2D:[8; 9] 99/291 (0.340)<br>height:[8; 10] 140/291 (0.481) |
| Different discovery GWAS, different ancestry | PTSD:[4] 248/1163 (0.213)<br>PTSD:[5] 227/1163 (0.195)<br>T2D:[6; 7] 241/1163 (0.207)<br>T2D:[7; 9] 248/1163 (0.213)<br>height:[8] 432/1163 (0.371)<br>height:[8; 10] 379/1163 (0.326) | PTSD:[4] 66/582 (0.113)<br>PTSD:[5] 65/582 (0.112)<br>T2D:[6; 7] 68/582 (0.117)<br>T2D:[7; 9] 66/582 (0.113)<br>height:[8] 155/582 (0.266)<br>height:[8; 10] 130/582 (0.223) | PTSD:[4] 24/291 (0.0825)<br>PTSD:[5] 17/291 (0.0584)<br>T2D:[6; 7] 19/291 (0.0653)<br>T2D:[7; 9] 12/291 (0.0412)<br>height:[8] 64/291 (0.220)<br>height:[8; 10] 55/291 (0.189) |

ABCD, Adolescent Brain and Cognitive Development Study; EUR, European-American ancestry; *n*, number of subjects; PTSD, post-traumatic stress disorder; T2D, type 2 diabetes.
Superscripts are the reference numbers for the discovery GWAS used to calculate PGS:
[4]Nievergelt et al. (2019), [5]Duncan et al. (2018), [6]Mahajan et al. (2018), [7]Chen et al. (2019), [8]Marouli et al. (2017), [9]Scott et al. (2017), [10]Wood et al. (2014).

| Table S14. LDSC estimates of SNP heritability for EUR GWAS: Observed vs. Liability Scale | | | | | |
|---|---|---|---|---|---|
| **Trait** | **Discovery GWAS** | **$h^2_{SNP}$ (SE) Observed Scale** | **$h^2_{SNP}$ (SE) Liability Scale** | **Sample[a] Prevalence** | **Population[b] Prevalence** |
| PTSD | Nievergelt et al. (2019)[4] | 0.0168 (0.003) | 0.0364 (0.0065) | 0.25 | 0.08 |
| | Duncan et al. (2018)[5] | 0.0966 (0.0427) | 0.1263 (0.059) | 0.13 | 0.08 |
| T2D | Scott et al. (2017)[9] | 0.0785 (0.0054) | 0.1431 (0.0099) | 0.16 | 0.08 |
| | Mahajan et al. (2018)[6] | 0.1945 (0.0083) | 0.3547 (0.0152) | 0.16 | 0.08 |
| Height | Marouli et al. (2017)[8] | 0.6967 (0.0674) | - | - | - |
| | Wood et al. (2014)[10] | 0.3119 (0.0141) | - | - | - |

LDSC, linkage disequilibrium score regression; EUR, European-American ancestry; PTSD, post-traumatic stress disorder; T2D, type 2 diabetes; $h^2_{SNP}$, SNP-based heritability; SE, standard error estimate obtained via block jackknife.

[a]Sample prevalence is the proportion of cases included in the discovery GWAS.

[b]We used the combined male and female population prevalence for PTSD that was cited by Duncan et al. (2018)[5] and the Centers for Disease Control and Prevention estimate of T2D prevalence for the population of white, non-Hispanic American adults[11] when calculating heritability on the liability scale.
